# DPP9 directly sequesters the NLRP1 C-terminus to repress inflammasome activation

**DOI:** 10.1101/2020.08.14.246132

**Authors:** L. Robert Hollingsworth, Humayun Sharif, Andrew R. Griswold, Pietro Fontana, Julian Mintseris, Kevin B. Dagbay, Joao A. Paulo, Steven P. Gygi, Daniel A. Bachovchin, Hao Wu

## Abstract

NLRP1 is a cytosolic inflammasome sensor that mediates activation of caspase-1, which in turn induces cytokine maturation and pyroptotic cell death^1-6^. Gain-of-function NLPR1 mutations cause skin inflammatory diseases including carcinoma, keratosis, and papillomatosis^7-14^. NLRP1 contains a unique function-to-find domain (FIIND) that autoproteolyzes into noncovalently associated subdomains^15-18^. Proteasomal degradation of the autoinhibitory N-terminal fragment (NT) activates NLRP1 by releasing the inflammatory C-terminal fragment (CT)^19,20^. Cytosolic dipeptidyl peptidases 8 and 9 (DPP8/9) interact with NLRP1, and small-molecule DPP8/9 inhibitors activate NLRP1 by poorly characterized mechanisms^11,19,21^. Here, we report cryo-EM structures of the human NLRP1-DPP9 complex, alone and in complex with the DPP8/9 inhibitor Val-boroPro (VbP). Surprisingly, the NLRP1-DPP9 complex is a ternary complex comprised of DPP9, one intact FIIND of a non-degraded full-length NLRP1 (NLRP1-FL) and one NLRP1-CT freed by NT degradation. The N-terminus of the NLRP1-CT unfolds and inserts into the DPP9 active site but is not cleaved by DPP9, and this binding is disrupted by VbP. Structure-based mutagenesis reveals that the binding of NLRP1-CT to DPP9 requires NLRP1-FL and vice versa, and inflammasome activation by ectopic NLRP1-CT expression is rescued by co-expressing autoproteolysis-deficient NLRP1-FL. Collectively, these data indicate that DPP9 functions as a “bomb-diffuser” to prevent NLRP1-CTs from inducing inflammation during homeostatic protein turnover.

## Main

The nucleotide-binding domain (NBD) and leucine-rich repeat (LRR)-containing (NLR) protein NLRP1 is the founding member of the NLR family that comprises most inflammasome sensors^1,16^. In addition to the FIIND, human NLRP1 also possesses an N-terminal pyrin domain (PYD) and a C-terminal caspase activation and recruitment domain (CARD), whereas rodent NLRP1 alleles do not contain PYDs^1,22^ (Fig. 1a). Recent studies demonstrated that functional N-terminal degradation by the proteasome is a mechanism for NLRP1 activation. *Bacillus anthracis* lethal factor and *Shigella flexneri* ubiquitin ligase IpaH7.8 directly cleave and ubiquitinate mouse NLRP1B, respectively, leading to proteasomal degradation of the autoinhibitory NLRP1-NT^19,20,23-26^. Because FIIND autoproteolysis leads to the break of the NLRP1 chain between the two FIIND subdomains ZU5 and UPA^15,17^, degradation of the NLRP1-NT releases noncovalently associated NLRP1-CT (or UPA-CARD) and facilitates CARD filament formation, recruitment of the apoptosis-associated speck-like protein containing a CARD (ASC) adaptor and pro-caspase-1, and inflammasome activation^19,20^. Active caspase-1 processes interleukin-1 (IL-1) family pro-cytokines to their bioactive forms, and cleaves gasdermin D (GSDMD) to liberate its pore-forming N-terminus^27-30^, which oligomerizes and perforates cell membranes to mediate cytokine secretion and inflammatory cell death^31-38^. Thus far, no such pathogen-derived activator of human NLRP1 or mouse NLRP1A has been identified^39^.

**Figure 1.**
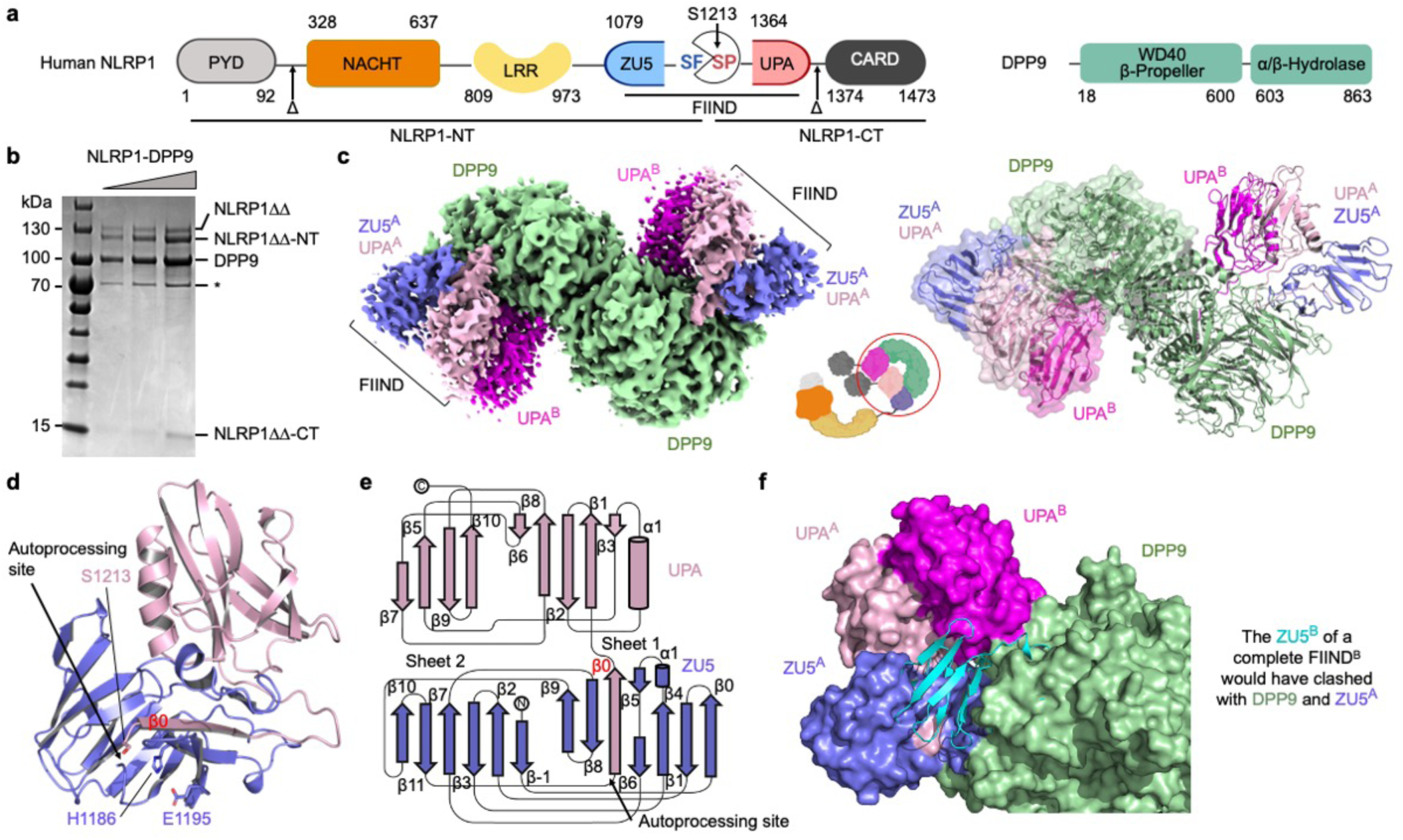
Structure of the NLRP1-DPP9 complex. **a**, Domain organization. **b**, SDS-PAGE of the purified NLRP1-DPP9 complex. HSP70 contamination is noted with an asterisk (*). **c**, Cryo-EM map (left) and the model (right) of the ternary NLRP1^A^-NLRP1^B^-DPP9 complex. The DPP9 dimer and the two copies of NLRP1 (A and B) are labelled with the colour scheme in (a). A schematic diagram (middle) denotes the entire NLRP1 and DPP9 molecules versus the ordered, resolved portions of the proteins (red circle). **d**, FIIND^A^ overview with ZU5 (blue) and UPA (light pink) subdomains. The catalytic triad residues (H1186, E1195, and S1213) are shown in sticks. **e**, Topology of the FIIND with secondary structures labelled. **f**, Superimposition of FIIND^A^ onto the UPA^B^. NLRP1^B^ has to be free NLRP1-CT because a ZU5 subdomain at site B would have clashed with ZU5 and UPA at site A and with DPP9.

In contrast to the selective activation of mouse NLRP1B by certain microbes, small-molecule dipeptidyl peptidase (DPP) inhibitors such as VbP (also known as Talabostat or PT-100) activate all functional NLRP1 alleles in human and rodents^19,21,39-41^. DPPs cleave substrates with a proline and, to a lesser extent, alanine, at their penultimate N-terminal amino acid position (NH_2_-X-P or NH_2_-X-A)^42^. Like pathogen-induced mouse NLRP1B activation, VbP-induced inflammasome activation relies on proteasomal degradation^11,19,43^. Two DPP family members, DPP8 and DPP9, directly bind NLRP1^11,43^, but how these DPPs suppress NLRP1 and how DPP inhibitors activate NLRP1 are unknown.

### Cryo-EM structure of the NLRP1-NLRP1-DPP9 ternary complex

Full-length human NLRP1 was prone to aggregation in multiple expression systems, and we identified PYD and CARD truncated NLRP1 (ΔΔ) with a C-terminal cleavable GFP-FLAG tag (NLRP1ΔΔ-TEV-GFP-FLAG) as a promising soluble construct when co-expressed with His-DPP9 in expi293F cells (Fig. 1b). Throughout the purification steps including anion exchange chromatography, both autoprocessed and unprocessed NLRP1 remained bound to DPP9 (Fig. 1b). We collected cryo-EM data on the NLRP1-DPP9 complex using a Titan Krios microscope equipped with a K3 Summit direct electron detector. Crosslinking with glutaraldehyde prior to plunging grids^44^ reduced complex dissociation, and we supplemented data collected at 0° with those at a stage tilt of 37° calculated by cryoEF^45^ to compensate for severe orientation preference. The final reconstructed map reached a resolution of 3.6 Å estimated by Fourier shell correlation of independent half maps (Extended Data Fig. 1). The atomic model was validated independently by crosslinking mass spectrometry; the ten intermolecular crosslinks identified by the amine-amine crosslinker bis(sulfosuccinimidyl)suberate (BS3) correspond to ∼10-20 Å Cα-Cα distances in the final model (Extended Data Fig. 2).

The cryo-EM structure surprisingly revealed that two NLRP1 molecules bind each monomer of the DPP9 dimer, forming an NLRP1^A^-NLRP1^B^-DPP9 ternary complex (Fig. 1c). The first NLRP1 (molecule A) is composed of a complete FIIND with intimately associated ZU5 and UPA subdomains (Fig. 1d-e, Extended Data Fig. 3a). The β-sandwich folds in these subdomains are similar to previously defined ZU5 and UPA domains^46,47^, but with a number of differences, in particular the insertion of the first β-strand (β0) of UPA into the ZU5 fold as if it is the last β-strand of ZU5 (Fig. 1d-e). Since our sample contained more autoprocessed than unprocessed NLRP1 (Fig. 1a), we built the model as processed FIIND. In this model of post-autoprocessed FIIND, the catalytic triad residues (S1213 of UPA, H1186 and E1195 of ZU5) are positioned nearly correctly for catalysis (Fig. 1d), suggesting that there are likely limited conformational changes between the unprocessed and processed forms.

The second NLRP1 (molecule B) contains only the UPA subdomain (Fig. 1c, Extended Data Fig. 3a). If we place the ZU5 subdomain of the second NLRP1 using the ZU5-UPA structure of the first NLRP1, it clashes with both DPP9 and the first NLRP1 (Fig. 1f), suggesting that the second NLRP1 cannot include the ZU5 subdomain or the N-terminal fragment. Thus, the NLRP1^A^-NLRP1^B^-DPP9 complex can only be formed when full-length NLRP1 and N-terminally degraded NLRP1 are both present. NACHT and LRR domains of NLRP1 are not visible in the density, likely due to a flexible linkage between the FIIND and the preceding LRR (Fig. 1a).

### NLRP1-CT inserts into the DPP9 substrate tunnel and is displaced by VbP

A striking observation in the NLRP1-DPP9 complex is that the N-terminal segment of UPA^B^ (β0, residue S1213-N1224) unfolds and inserts into the DPP9 substrate tunnel as an extended polypeptide chain (Fig. 2a). The inserted segment in UPA^B^ shows dramatically different conformation from its UPA^A^ counterpart, whereas the gross conformations of UPA^A^ and UPA^B^ are similar (Extended Data Fig. 3b-c). The terminal amino group of the segment forms hydrogen-bond and salt-bridge interactions with E248 and E249 in the EE-helix of DPP9, in a manner that mimics substrate binding^48^. The DPP9 active site is also known to undergo substantial rearrangement at a large loop segment, which partially folds into an α-helix known as the R-helix when an Arg residue on the helix engages with a substrate^48^. In the NLRP1-DPP9 complex, R133 of DPP9 interacts with both E248 in the EE-helix and main chain carbonyl oxygen atoms in the N-terminal segment of UPA^B^, and the R-helix undergoes a disorder to order transition (Fig. 2a), similar to substrate binding. However, we show below that UPA^B^ binds in a modified manner to avoid being cleaved by DPP9.

**Figure 2.**
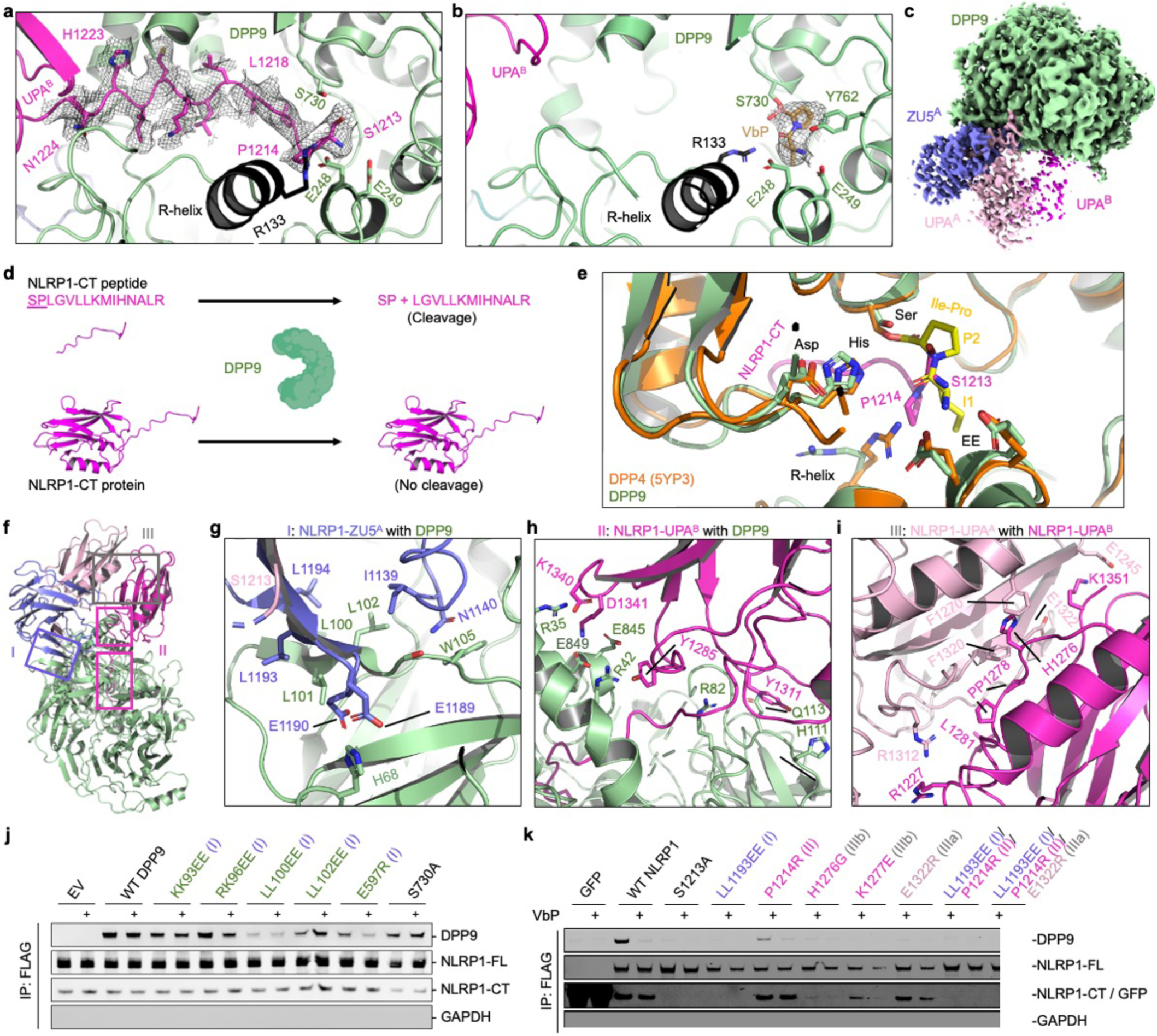
Detailed interfaces in the NLRP1-DPP9 ternary complex and inhibition by the DPP9 inhibitor VbP. **a**, Insertion of the N-terminal peptide of UPA^B^ into the DPP9 active site. **b**, Displacement of the UPA^B^ N-terminal peptide from the DPP9 active site by VbP. **c**, Cryo-EM map of the NLRP1-DPP9 complex in the presence of VbP. VbP binding reduces UPA^B^ occupancy. **d**, Ability of DPP9 to cleave an isolated UPA N-terminal peptide, but not an entire UPA. **e**, Comparison of the binding modes of the UPA^B^ N-terminal peptide in the NLRP1-DPP9 complex and the Ile-Pro dipeptide in an acyl-enzyme intermediate^49^. **f**, Overview of three interfaces important for NLRP1-DPP9 association. Regions blocked in rectangles are shown in detail in (a) and (g-i). **g-i**, Zoom-ins of each NLRP1-DPP9 binding interface. **j**, FLAG co-immunoprecipitation using FLAG-tagged WT NLRP1 and His-tagged WT and mutant DPP9. Anti-DPP9, anti-NLRP1-CT (NLRP1-FL and NLRP1-CT), and anti-GAPDH antibodies were used in the immunoblots. EV: empty vector. The Roman numerals in the parentheses represent the three interfaces in the NLRP1-DPP9 ternary complex. Each immunoblot is representative of > 2 independent experiments. **k**, FLAG co-immunoprecipitation using FLAG-tagged WT or mutant NLRP1 and His-tagged WT DPP9. FLAG-tagged GFP was used as a negative control. Anti-DPP9, anti-FLAG (NLRP1-FL, NLRP1-CT, and GFP), and anti-GAPDH antibodies were used in the immunoblots. The Roman numerals in parentheses represent the three interfaces in the NLRP1-DPP9 ternary complex. Each immunoblot is representative of > 2 independent experiments.

Since small-molecule DPP8/9 inhibitors also bind their active sites^48^ and in turn activate NLRP1^41^, we explored what structural effects they would have on the NLRP1-DPP9 complex. To this end, we solved a 2.9 Å cryo-EM structure of the NLRP1-DPP9 complex purified in the presence of the small-molecule inhibitor VbP, a covalent peptidomimetic that modifies the catalytic S730 (Fig. 2b, Extended Data Fig. 4, 5a). The density of VbP facilitated its unambiguous placement, which showed that VbP interacts with several notable DPP9 residues in addition to the catalytic S730: the R-loop residue R133 and the EE motif composed of E248 and E249 (Fib. 2b, Extended Data Fig. 5b). VbP also induces the substrate-bound state of DPP9 as indicated by ordering of the R-helix. In fact, the VbP conformation closely resembles the pose of an isoleucine-proline substrate bound to bacterial DPP4^49^ (Extended Data Fig. 5c).

Remarkably, VbP completely displaced the NLRP1 UPA^B^ peptide from the DPP9 substrate cavity (Fig. 2b). Additionally, the cryo-EM density of the entire UPA^B^ molecule in the NLRP1-DPP9-VbP complex structure was notably weaker (Fig. 2c), suggesting that by removing key stabilizing interactions in the substrate tunnel, VbP greatly reduces the affinity of UPA^B^ of NLRP1-CT to DPP9 or might even displace UPA^B^ entirely. These structural data rationalize prior reports that VbP weakened the NLRP1-DPP9 interaction as shown by co-immunoprecipitation and mass spectrometry^11,43^.

### The DPP9 substrate sequence of NLRP1-CT cannot be cleaved by DPP9

NLRP1-CT contains a proline in the P2 position (NH_2_-S-P) (Fig. 1a). Since this N-terminus threads into the active site tunnel of DPP9 (Fig. 2a), we wondered if DPP9 could cleave its first two residues. We performed Edman degradation on the human NLRP1-CT band from the purified NLRP1-DPP9 complex (Fig. 1b), which showed no cleavage by DPP9 (Fig. 2d, Extended Data Fig. 6a). We also performed an experiment of chemical enrichment of protease substrates (CHOPS), which uses a 2-pyridinecarboxaldehyde (2PCA)-biotin probe to selectively biotinylate free N-termini except those with proline in the second position^42^. Lysates from HEK293T cells doubly deficient in *DPP8 and DPP9* and ectopically expressing NLRP1 were treated with recombinant DPP9 prior to performing CHOPS. DPP9 treatment did not increase NLRP1-CT signal, indicating that NLRP1-CT was not cleaved by DPP9 (Fig. 2d, Extended Data Fig. 6b).

In contrast, a peptide comprising the first 15-residues of NLRP1-CT is a bona fide DPP9 substrate shown by mass spectrometry analysis of the reaction product (Fig. 2d, Extended Data Fig. 6c) and by its inhibition of DPP9 activity against the fluorogenic Ala-Pro-7-amino-4-methylcoumarin (AP-AMC) substrate (Fig. 2d, Extended Data Fig. 6d). Therefore, the N-terminal substrate motif must have been constrained in the context of NLRP1-CT so that no cleavage is possible. Consistent with this hypothesis, the conformation of the NLRP1-CT N-terminal substrate motif bound at the DPP9 active site is markedly different from that of the isoleucine-proline substrate bound at bacterial DPP4 active site (PDB ID: 5YP3)^49^; NLRP1-CT situates further back in the DPP9 tunnel (Fig. 2e). These data suggest that although NLRP1-CT has a DPP9 substrate sequence, it uses a modified binding mode to avoid cleavage (Fig. 2d). Thus, regulation of NLRP1 by DPP9 does not involve DPP9-mediated cleavage of NLRP1-CT.

### Three interfaces mediate the NLRP1-NLRP1-DPP9 ternary complex assembly

There are three interaction sites in the NLRP1^A^-DPP9-NLRP1^B^ complex: interface I (between ZU5^A^ and DPP9), interface II (between UPA^B^ and DPP9), and interface III (between UPA^A^ and UPA^B^) (Fig. 2f). UPA^A^ has limited direct contact with DPP9; however, it likely helps to recruit the second NLRP1 via interface III. All three interfaces are extensive, burying ∼1,000 Å^2^, ∼1,600 Å^2^ and ∼800 Å^2^ surface areas per partner, respectively.

Interface I is formed by regions of β9 and *α*1-β5 loop in ZU5^A^ and the WD40 domain in DPP9, with part of the interactions through hydrogen bonds between β-strands (Fig. 2g). Residues L1193, L1194 (β9), I1139 and N1140 (*α*1-β5) in ZU5^A^ pack against L100, L101, L102, S104, W105 (WD40 blade I) and E597 (WD40 blade 8) in DPP9. E1189 and E1190 (β9-β10 loop) of ZU5^A^ interact with H68 (WD40 blade I) and a main chain amide in DPP9. Part of the blade I of DPP9 is disordered in the DPP9 crystal structure^48^ but becomes ordered upon NLRP1 interaction (Extended Data Fig. 7a). The β8 and β9 strands also contain the FIIND catalytic residues histidine (H1186) and glutamic acid (E1195), while the catalytic serine (S1213) is the first residue of the UPA subdomain (Fig. 1a), which may explain the sensitivity of autoprocesing to interface I mutations (below).

Interface II has two components, one formed by the N-terminal segment of UPA^B^ (S1213-N1224) as described previously (Fig. 2a), and the other formed between one face of the UPA^B^ globular domain and the α/β-hydrolase domain of DPP9 (Fig. 2h). The latter includes residues Y1285 (β5-β6), Y1311 (β7-β8), K1340, and D1341 (β9-β10) in UPA^B^ and R35, R42 (*α*1), R82 (blade I), P845, and E846 (*α*12) in DPP9.

In interface III, UPA^A^ interacts intimately with UPA^B^ using the sides of their β-sandwich folds (Fig. 1d-e, 2i), comprising both hydrogen bonding interactions and hydrophobic packing between the two UPAs. The relationship between these two subdomains is mainly a translation with only a 9° rotation between them (Extended Data Fig. 3b), and this UPA-UPA interaction has never been observed in other ZU5 and UPA-containing structural homologs^46,47^. Given the translational relationship between these UPAs, the entire interaction surface differs between UPA^A^ (IIIa) and UPA^B^ (IIIb). A few residues are of note, including D1317, L1319 and E1322 (β8) of UPA^A^, and H1276, K1277, P1278 and P1279 (β3-β5 loop) of UPA^B^.

### Structure-based NLRP1 and DPP9 mutations compromise complex formation

We generated structure-based mutations on all three interfaces to determine their effects on the NLRP1-DPP9 complex formation. NLRP1 interface I mutations were remarkably prone to disrupting FIIND autocatalytic activity. Therefore, we used reciprocal mutants on DPP9, expressed in *DPP8/9* knockout (KO) cells (Extended Data Fig. 7b), to validate the importance of this interface. Both LL100EE and E597R DPP9 mutants strongly reduced NLRP1 interaction (Fig. 2j) without compromising DPP9’s post-proline cleavage activity or sensitivity to VbP (Extended Data Fig. 7c-d). We also managed to obtain the DPP9-binding defective NLRP1 interface I mutant LL1193EE, which although showed a defect in FIIND autoprocessing, is autoactive without VbP (Fig. 2k, 3a, Extended Data Fig. 7e), suggesting that it retained at least some autocatalytic activity. The interface II mutant P1214R that resides at the N-terminus of the UPA, which is also a previously described autoactive patient mutation^11,12^, was fully autoprocessed but reduced DPP9 binding (Fig. 2k, Extended Data Fig. 7e). As expected, addition of VbP during co-IP further reduced the bound DPP9 or bound NLRP1 (Fig. 2j-k). Mutations on interface III, the UPA^A^ (IIIa)-UPA^B^ (IIIb) dimerization surface, almost completely abolished DPP9 binding while retaining autoprocessing (Fig. 2k, Extended Data Fig. 7e). Similarly, the S1213A autoprocessing mutation, or wild-type (WT) NLRP1 treated with VbP, also abolished DPP9 binding in the co-IP assay. These data suggest that there is cooperativity between the two NLRP1 molecules in binding DPP9, in which neither NLRP1^A^ nor NLRP1^B^ binds to DPP9 without the other. In vitro however, the high protein concentration could facilitate the binding of NLRP1^A^ to DPP9 even when NLRP1^B^ is largely absent (Fig. 2c).

### Loss of DPP9 interaction causes NLRP1 inflammasome autoactivation and UPA-UPA interaction is required for both NLRP1 inhibition and activation

Since small-molecule DPP9 inhibition and its genetic ablation activates NLRP1^11,39,41^, we hypothesized that loss of DPP9 interaction would result in NLRP1 autoactivation. We designed a HEK293T system that recapitulates key components of the inflammasome pathway to evaluate the functional consequences of specifically disrupting interactions of NLRP1-FL^A^, NLRP1-CT^B^, or both with DPP9. Specifically, we transiently reconstituted ASC and loss-of-DPP9 interaction NLRP1 mutations in HEK293T cells stably expressing caspase-1 and GSDMD, and assayed cell death by release of the mitochondrial enzyme lactate dehydrogenase (LDH). Indeed, mutations on interfaces I and II, including the previously reported patient mutation P1214R^11,12^, caused marked cell death and GSDMD processing in the absence of VbP, with no noticeable further increase upon VbP addition (Fig. 3a). Concomitantly, these NLRP1 mutants spontaneously formed ASC specks (Fig. 3b, Extended Data Fig. 8a-b) in the absence of any activating signal. Thus, disruption of either NLRP1-DPP9 interface I or II leads to spontaneous inflammasome activation, likely mediated by unrestrained NLRP1-CT oligomerization, ASC recruitment, and downstream signalling.

**Figure 3.**
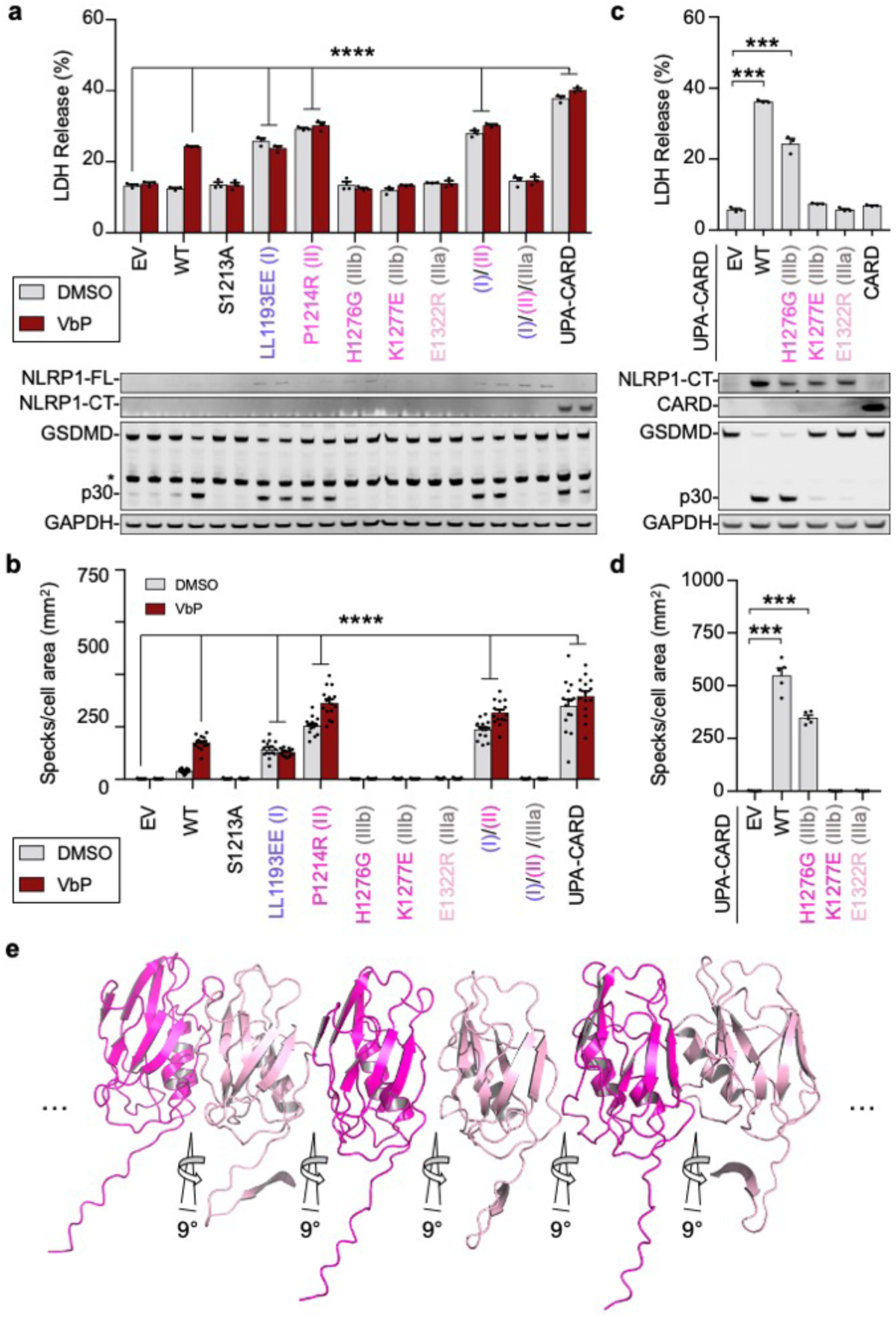
Functional consequences of interfacial mutations in the NLRP1-DPP9 ternary complex. Interface I and II mutants cause NLRP1 autoactivation and UPA-UPA interactions are required for inflammasome activity. **a**, LDH release (top) and GSDMD processing (bottom) from transient expression of empty vector (EV), WT or mutant NLRP1-FL, or UPA-CARD in a reconstituted HEK293 inflammasome system, with and without addition of VbP. Anti-NLRP1-CT (FL and CT), anti-GSDMD, and anti-GAPDH antibodies were used in the immunoblots. *: non-specific bands; p30: GSDND N-terminal fragment from caspase-1 cleavage. **b**, Quantification of speck formation induced by expression of EV, WT or mutant NLRP1-FL, or UPA-CARD in the presence and absence of VbP. **c**, LDH release (top) and GSDMD processing (bottom) by direct expression of WT or mutant NLRP1 UPA-CARDs. CARD alone was also included. **d**, Quantification of speck formation induced by expression of WT or mutant NLRP1 UPA-CARD. **e**, A modelled UPA oligomer based on the near front-to-back interaction in the NLRP1-DPP9 ternary complex. In the model, the N-terminal tails of free UPAs are shown in either the UPA^A^ or UPA^B^ conformation in complex with DPP9, but in reality, this conformation is likely to be different. All data are representative of > 3 independent experiments. **** in (a) and (b): p < 0.0001 compared to EV by 2-way ANOVA with Tukey multiple comparison correction. *** in (c) and (d): p < 0.001 by two-sided Student’s t test.

Surprisingly, interface III UPA^A^-UPA^B^ loss-of-interaction mutants were completely deficient in both spontaneous and VbP-mediated inflammasome signalling (Fig. 3a-b) despite autoprocessing at close to wild-type levels especially for K1277E (IIIb) and E1322R (IIIa) (Fig. 2k). Furthermore, addition of an interface III mutant abolished the activity of the interface I/II double mutant (Fig. 3a-b). We reasoned that interface III UPA-UPA contacts on the inflammasome are important for productive signalling, since the UPA itself lowers the threshold for productive CARD signaling^50,51^. To test this hypothesis, we ectopically expressed NLRP1-CT in the same reconstituted HEK293T cell system, which as expected induced cell death and ASC speck formation (Fig. 3c-d, Extended Data Fig. 8c-d). The interface III mutations K1277E and E1322R abolished CT-mediated cell death, GSDMD cleavage, and ASC speck formation, while H1276G reduced CT-mediated inflammasome activation (Fig. 3c-d, Extended Data Fig. 8c-d). An NLRP1-CARD construct was similarly deficient in inducing cell death (Fig. 3c).

This requirement for the UPA domain and the UPA-UPA interface III contacts in NLRP1-CT inflammasome activation suggests that UPA may play an architectural role in inflammasome assembly. Interestingly, the UPA domain within an intact FIIND cannot self-oligomerize due to clashes at the ZU5 domain (Fig. 1f, Extended Data Fig. 9a). In addition, freed NLRP1-CT cannot be further recruited to the ternary complex by an UPA-UPA interaction with either UPA^A^ due to a clash with ZU5^A^ or with UPA^B^ due to clashes with both monomers of the DPP9 dimer (Extended Data Fig. 9b). However, the UPA domains in freed NLRP1-CTs could in principle oligomerize assuming that they maintain the same near front-to-back binding mode seen in the complex with DPP9 (Fig. 3e). This UPA oligomerization would create a platform for CARD oligomerization, ASC recruitment, and inflammasome assembly. Thus, the UPA domain has two opposing functions: to mediate repression of the NLRP1-CT through DPP9 association and to facilitate its activation by promoting oligomerization.

### DPP9 requires NLRP1-FL to sequester NLRP1-CT

Because mutations on either side of the UPA^A^-UPA^B^ interface almost completely disrupted NLRP1 association with DPP9 while retaining autoprocessing (Fig. 2i), we reasoned that DPP9 binding by NLRP1-FL and NLRP1-CT is cooperative and that NLRP1-FL might dimerize with NLRP1-CT. We co-expressed FLAG-tagged full-length NLRP1 S1213A that is autoproteolytically deficient and can only occupy site A with a construct for NLRP1-CT that can only occupy site B (Fig. 4a). For NLRP1-CT expression, we used an N-terminal ubiquitin (Ub) fusion (Ub-NLRP1-CT), which gets co- or post-translationally processed^52^ to generate the native S1213 N-terminus. Indeed, NLRP1-S1213A captured NLRP1-CT, and co-expression of these two components rescued DPP9 association (Fig. 4b), confirming directly that DPP9 engagement is cooperative.

**Figure 4.**
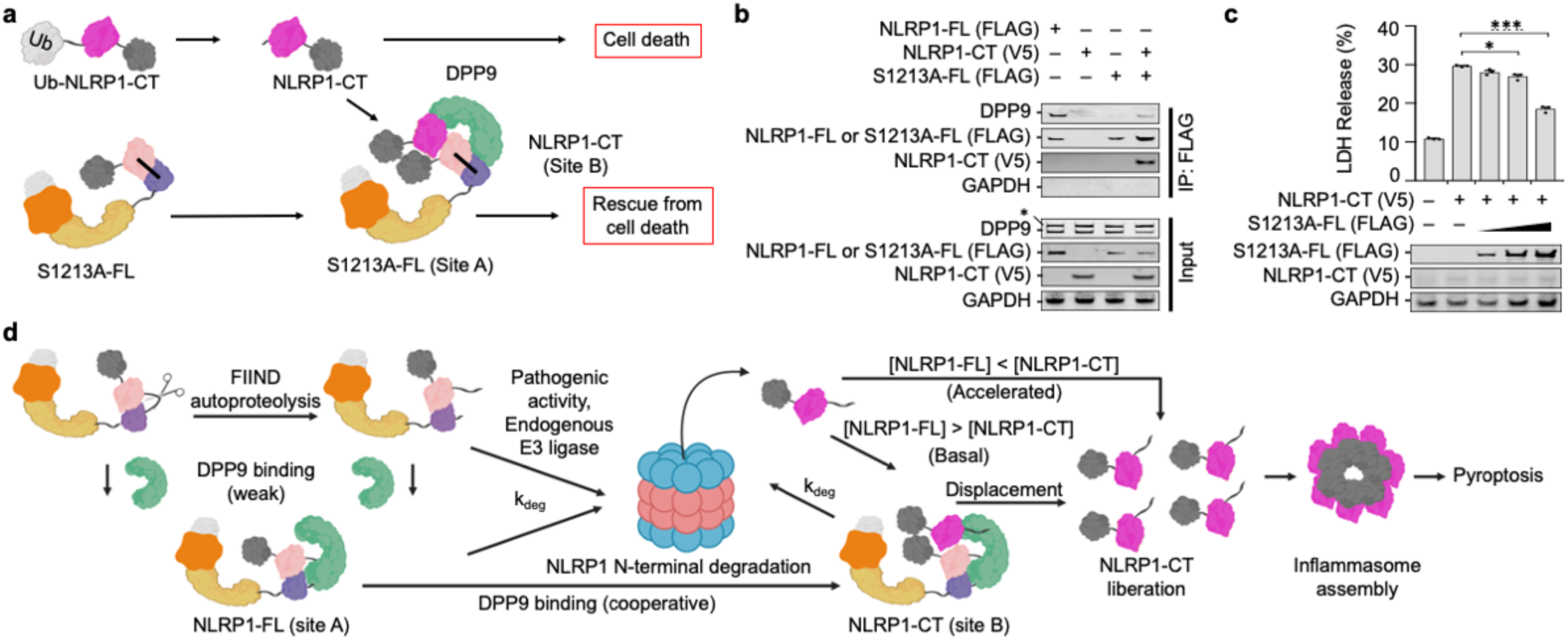
Repression of NLRP1-CT inflammasome activity by tertiary complex formation. **a**, Schematic representation of tertiary complex formation between the autocatalysis-deficient NLRP1-S1213A-FL, Ub-NLRP1-CT, and DPP9. Ub-NLRP1-CT is co- or post-translationally processed by endogenous deubiquitinases, resulting in NLRP1-CT with the native S1213 N-terminus. **b**, FLAG co-immunoprecipitation using FLAG-tagged WT NLRP1-FL or S1213A-FL and V5-tagged Ub-NLRP1-CT. Neither S1213A-FL (site A) nor NLRP1-CT (site B) alone pulled down endogenous DPP9. In contrast, co-expression of the two constructs formed a ternary complex with DPP9. **c**, Suppressed NLRP1-CT-induced cell death measured by LDH release in a reconstituted HEK293 inflammasome system when co-expressed with increasing amount of S1213A-FL. All data are representative of > 2 independent experiments. *: p < 0.05 and ***: p < 0.001 by two-sided Student’s t test. **d**, Schematic of NLRP1 activation and repression by DPP9. NLRP1-FL, together with DPP9, represses some threshold of free NLRP1-CTs. Enhanced degradation of NLRP1-FL, or displacement of the NLRP1-CT, leads to inflammasome signalling.

Because DPP9-engagement of NLRP1-CT into the tertiary complex should prevent UPA-UPA oligomerization (Extended Data Fig. 9b), and inhibit inflammasome activity, we tested this hypothesis by co-expressing NLRP1-S1213A-FL with Ub-NLRP1-CT in the reconstituted HEK system. Indeed, NLRP1-S1213A-FL titration reduced NLRP1-CT-mediated cell death in a dose-dependent manner (Fig. 4c). Thus, DPP9 quenches the inflammatory activity of NLRP1-CT in a manner that depends on the presence of NLRP1-FL.

## Conclusion

In summary, our structural and cellular data show that DPP9, together with NLRP1-FL, acts as a checkpoint on NLRP1 inflammasome activation by directly binding to and sequestering the inflammatory NLRP1-CT. We speculate that free inflammatory NLRP1-CTs are likely generated by some as-yet-unknown danger-associated signal, but also during normal homeostatic protein turnover. DPP9 and NLRP1-FL binding would therefore act to avoid spurious inflammation from low, background levels of NLRP1-CT relative to NLRP1-FL. However, significant and rapid increases in the ratio of NLRP1-CT to NLRP1-FL would overwhelm this checkpoint and lead to inflammasome activation and pyroptotic cell death. Our data also show how DPP8/9 inhibitors and the patient-derived P1214R germline mutation^53^ interfere with the binding of the NLRP1-CT to DPP9, and thereby attenuate the ability of DPP9 and NLRP1-FL to sequester these inflammatory fragments. Interestingly, it should be noted that inhibition of DPP8/9’s enzymatic activity also appears to induce NLRP1 N-terminal degradation^19,41^, and thus DPP8/9 inhibitors likely promote NLRP1 inflammasome activation via two separate mechanisms (Fig. 4d). Overall, we expect that future studies based on these results will not only lead to greater mechanistic understanding of this long enigmatic inflammasome, but also eventually enable therapeutic modulation of NLRP1 to treat human disease.

## Methods

### Constructs and cloning

Mammalian codon-optimized full-length human NLRP1 cDNA (isoform 1, Uniprot ID Q9C000-1) was synthesized by Synbio Technologies. Codon-optimized NLRP1 lacking the PYD and CARD (Residues W148-P1364, NLRP1ΔΔ) was subcloned into an in-house modified pcDNA3.1 LIC 6D (Addgene plasmid #30127) vector containing a C-terminal TEV linker, GFP tag, and FLAG tag (NLRP1ΔΔ-TEV-GFP-FLAG). NLRP1-FL was also cloned into pcDNA3.1 LIC 6A (Addgene plasmid #30124) with a C-terminal FLAG tag (NLRP1-FLAG). The short isoform of DPP9 (DPP9S, Uniprot ID Q86TI2-1) from the Harvard PlasmID Database was subcloned into pfastBac HTB (N-terminal His-TEV tag) for recombinant protein expression. DPP9 was then subcloned into pcDNA3.1 LIC 6A with either an N-terminal His- or FLAG-tag (His-TEV-DPP9 or FLAG-TEV-DPP9). Point mutations were introduced with Q5 site-directed mutagenesis (NEB) or QuickChange site-directed mutagenesis kit (Agilent). Ub-fused NLRP1 constructs were synthesized (GenScript) with an N-terminal synthetic ubiquitin sequence followed by NLRP1-CT (S1213-S1473) and cloned into the pcDNA3.1 vector.

### Protein expression, purification, and Edman degradation

Recombinant DPP9 was purified similarly to Ross *et al*., 2018^48^. Baculoviruses containing DPP9 were prepared using the Bac-to-Bac system (Invitrogen), and used to generate baculovirus-infected insect cells. To express DPP9, 1 mL of these baculovirus-containing cells was used to infect each L of Sf9 cells. Cells were harvested 48 h after infection by centrifugation (2,250 RPM, 20 min), washed once with phosphate buffered saline (PBS), flash-frozen in liquid nitrogen, and stored at -80 °C. The thawed pellet from 2 L of cells was resuspended in lysis buffer (80 mL, 25 mM Tris-HCl pH 8.0, 150 mM NaCl, 1 mM tris(2-carboxyethyl)phosphine abbreviated as TCEP, 5 mM imidazole), sonicated (3 s on 5 s off, 3.5 min total on, 45% power, Branson), and centrifuged at 40,000 RPM for 1.5 h (45 Ti fixed-angle rotor, Beckman). After centrifugation, the supernatant was incubated with 1 mL Ni-NTA resin at 4 °C for 1 h. The bound Ni-NTA beads washed once in batch and subsequently by gravity flow using 50-100 CV wash buffer (25 mM Tris-HCl pH 8.0, 150 mM NaCl, 1 mM TCEP, 25 mM imidazole). The protein was eluted with buffer containing 500 mM imidazole (5 mL), spin concentrated to 0.5 mL (Amicon Ultra, 100 kDa MW cutoff), and further purified by size exclusion chromatography (25 mM Tris-HCl pH 7.5, 150 mM NaCl, 1 mM TCEP) on a Superdex 200 increase 10/300 GL column (Cytiva). The yield was ∼5 mg per L of insect cell culture.

To purify the NLRP1-DPP9 complex, expi293F cells (1 L, 2-3×10^6^ cells/mL) were co-transfected with NLRP1ΔΔ-TEV-GFP-FLAG (0.7 mg) and the short isoform of DPP9 (His-TEV-DPP9, 0.3 mg) following incubation with polyethylenimine (3 mL, 1 mg/mL) in OptiMEM (100 mL) for 30 min. 24 h later, cells were supplemented with glucose (9 mL, 45%) and valproic acid (10 mL, 300 mM). Cells were harvested 5 d after transfection by centrifugation (2,000 RPM, 20 min), washed once with PBS, flash-frozen in liquid nitrogen, and stored at -80 °C. Later, the thawed pellet from 1 L of cells was resuspended in lysis buffer (50-100 mL, 25 mM Tris-HCl pH 7.5, 150 mM NaCl, 1 mM TCEP), sonicated (2 s on 8 s off, 3.5 min total on, 40% power, Branson), and ultracentrifuged at 40,000 RPM for 1 h (45 Ti fixed-angle rotor, Beckman). The soluble proteome was incubated with pre-equilibrated anti-FLAG M2 affinity gel (Sigma, 1.5 mL) for 4 h at 4 °C, washed in batch once with 5-10 column volume (CV) lysis buffer, and then washed by gravity flow with 25-50 CV lysis buffer. The NLRP1-DPP9 complex was eluted by on-column TEV protease cleavage at room temperature for 1 h using elution buffer (5 mL, 25 mM Tris-HCl pH 7.5, 150 mM NaCl, 5 mM MgCl_2_, 0.2 mM ADP, 1 mM TCEP, 0.2 mg TEV protease) and loaded onto a Mono Q 5/50 GL anion exchange column (Cytiva). Protein was eluted using a salt gradient from 150 mM to 1 M NaCl (25 mM Tris-HCl pH 8.0, 1 mM TCEP) over 15 CV. Subsequent purification steps employing size exclusion columns led to complete loss of protein, likely due to hydrophobic interactions between the protein and the column’s support. Nonetheless, the anion exchange eluent showed sufficiently homogeneous NLRP1-DPP9 complexes by negative stain electron microscopy. Mono Q eluent containing the complex was concentrated to 0.3-0.5 mg/mL (assuming ε = 1.25) using a 0.5 mL spin concentrator (Amicon Ultra, 50 kDa MW cutoff) and immediately supplemented with 0.2 mM ADP and 5 mM MgCl_2_. Concentrated eluent was dialyzed overnight into a HEPES buffer that is compatible with crosslinking for cryo-EM (25 mM HEPES pH 7.5, 150 mM NaCl, 5 mM MgCl_2_, 0.2 mM ADP, 1 mM TCEP) using a 0.5 mL Slide-A-Lyzer (ThermoFisher). Total protein yield varied between 0.2-0.5 mg per L of mammalian culture. Purified protein was run on an SDS-PAGE gel, transferred to a PVDF membrane (iBLOT 2), and stained by Coomassie Brilliant Blue. The NLRP1-CT band was excised and sent to the Tufts University Core Facility for Edman Degradation.

Expression and purification of the NLRP1-DPP9-VbP complex was identical to NLRP1-DPP9 (described above) except for the addition of 10 µM VbP to all purification and dialysis buffers. Extended protocol including intermediary purification and quality control results will be made available on protocols.io (https://www.protocols.io/groups/hao-wu-lab).

### Crosslinking mass spectrometry

Amine-amine crosslinking of the purified NLRP1-DPP9 complex was performed at a final concentration of 0.24 mg/mL using either 0.5, 1, or 2 mM bis(sulfosuccinimidyl)suberate (BS3) in 50 mM HEPES pH 7.8 and 100 mM NaCl for 1 h at room temperature. The reaction was quenched with hydroxylamine to a final concentration of 100 mM. Urea was added to 5 M concentration and samples were reduced for 1 h in 10 mM TCEP and 50 mM HEPBS pH 8.3, followed by alkylation with 30 mM iodoacetamide in the dark for one hour and quenching with 50 mM β-mercaptoethanol. Samples were then diluted with 50 mM HEPBS pH 8.3 to reduce urea concentration down to 1 M and digested with trypsin (Promega) at 1:10 enzyme:substrate ratio overnight at 37°C. Digested peptides were acidified with 10% formic acid (FA) to pH ∼2 and desalted using stage tips with Empore C18 SPE Extraction Disks (3M) and dried under vacuum.

Crosslinked samples were reconstituted in 5% FA/5% acetonitrile (ACN) and analysed in the Orbitrap Fusion Lumos Mass Spectrometer (ThermoFisher) coupled to an EASY-nLC 1200 (ThermoFisher) ultra-high pressure liquid chromatography (UHPLC) pump, as well as a high-field asymmetric waveform ion mobility spectrometry (FAIMS) FAIMSpro interface. Peptides were separated on an in-house column with 100 μM inner diameter and packed with 35 cm of Accucore™ C18 resin (2.6 μm, 150 Å, ThermoFisher), using a gradient consisting of 5–35% (ACN, 0.125% FA) over 135 min at ∼500 nL/min. The instrument was operated in data-dependent mode. FTMS1 spectra were collected at a resolution of 120,000, with an automated gain control (AGC) target of 5 × 10^5^, and a max injection time of 50 ms. The most intense ions were selected for MS/MS for 1 s in top-speed mode, while switching among three FAIMS compensation voltages: −40, –60, and –80 V in the same method. Precursors were filtered according to charge state (allowed 3 <= z <= 7), and monoisotopic peak assignment was turned on. Previously interrogated precursors were excluded using a dynamic exclusion window (60 s ± 7 ppm). MS2 precursors were isolated with a quadrupole mass filter set to a width of 0.7 Th and analysed by FTMS2, with the Orbitrap operating at 30,000 resolution, an AGC target of 100,000, and a maximum injection time of 150 ms. Precursors were then fragmented by high-energy collision dissociation (HCD) at a 32% normalized collision energy.

Mass spectra were processed and searched using the PIXL search engine^54^. Precursor tolerance was set to 15 ppm and fragment ion tolerance to 10 ppm. Methionine oxidation was set as a variable modification in addition to mono-linked mass of +156.0786 for BS3. Crosslinker mass shift of +138.0681 was used for BS3 reagent. All crosslinked searches included 50 most abundant protein sequences to ensure sufficient statistics for false discovery rate (FDR) estimation. Matches were filtered to 1% FDR on the unique peptide level using linear discriminant features as previously described^54^.

### Cryo-EM grid preparation and data collection

The purified NLRP1-DPP9 complex (0.3-0.5 mg/mL, 25 mM HEPES pH 7.5, 150 mM NaCl, 5 mM MgCl_2_, 0.2 mM ADP, 1 mM TCEP, ±10 μM VbP) was crosslinked with 0.02% glutaraldehyde on ice for 5 min and immediately loaded onto a glow-discharged Quantifoil grid (R1.2/1.3 400-mesh gold-supported holey carbon, Electron Microscopy Sciences), blotted for 3–5 s under 100% humidity at 4 °C, and plunged into liquid ethane using a Mark IV Vitrobot (ThermoFisher). These sample conditions were optimized extensively using insights gleaned from different small datasets collected at the Pacific Northwest Center for Cryo-EM (PNCC), the NCI’s National Cryo-EM Microscope Facility (NCEF), and the Harvard Cryo-EM Center (HMS). Final collection parameters are summarized in Extended Data Table 1.

Prior to data collection, all grids were screened for ice and particle quality. For data collection, movies were acquired at HMS using a Titan Krios microscope (ThermoFisher) at an acceleration voltage of 300 keV equipped with BioQuantum K3 Imaging Filter (Gatan, slit width 20 eV). Movies were recorded with a K3 Summit direct electron detector (Gatan) operating in counting mode at 105,000 x magnification (0.825 Å/pix).

For the NLRP1-DPP9 data set at 0° stage tilt: 7,840 movies were collected using SerialEM^55^ to vary the defocus range between −1 to −2.5 μm and to record two shots for each of the four holes per stage movement through image shift. All movies were exposed with a total dose of 67.5 e^-^/Å^2^ for 1.8 s fractionated over 45 frames. An ideal tilt angle of 37° was calculated from this processed dataset using cryoEF^56^. For the NLRP1-DPP9 data set at a stage tilt of 37°: 1,916 movies were collected using SerialEM^55^ to vary the defocus range between −1.5 to −3.0 μm with two shots recorded for each hole per stage movement through image shift. All movies were exposed with a total dose of 67.6 e^-^/Å^2^ for 2.6 s fractionated over 45 frames.

For the NLRP1-DPP9-VbP data set at 0° stage tilt: 3,553 movies were collected using SerialEM^55^ to vary the defocus range between −0.8 to −2.2 μm and to record two shots for each of the four holes per stage movement through image shift. All movies were exposed with a total dose of 52.0 e^-^/Å^2^ for 2.0 s fractionated over 50 frames. An ideal tilt angle of 37° was calculated from this processed dataset using cryoEF^56^. For NLRP1-DPP9-VbP at a stage tilt of 37°: 1,954 movies were collected using SerialEM^55^ to vary the defocus range between −1.5 to −3.0 μm with two shots recorded for each hole per stage movement through image shift. All movies were exposed with a total dose of 65.0 e^-^/Å^2^ for 2.4 s fractionated over 43 frames.

### Cryo-EM data processing

Data processing leveraged SBgrid Consortium^57^ and NSF XSEDE^58^ software and computing support (Extended Data Fig. 1, 4). Movies collected at HMS were pre-processed on-the-fly by the facility’s pipeline script. Movies were corrected by gain reference and for beam-induced motion, and summed into motion-corrected images using the Relion 3.08 implementation of the MotionCor2 algorithm^59^. CTFFIND4^60^ or goCTF^61^ was used to determine the defocus of each micrograph (Extended Data Fig. 1, 4). These pre-processed micrographs were used for subsequent analysis.

For the NLRP1-DPP9 complex collected at 0° stage tilt, crYOLO^62^ (generalized training by the HMS facility) picked 1,1197,166 particles from the 7840 micrographs. Relion 3.1^63^ was used for subsequent image processing. Initially, picked particles were subjected to multiple rounds of 2D classification until the classes appeared visually homogenous. A randomized set of 100,000 particles was selected for the *de novo* reconstruction of an initial model, which was low-pass-filtered to 40 Å to use as the input reference map for 3D classification. Multiple rounds of 3D classification were used to produce homogeneous particle stacks among the heterogeneous particle populations (unbound DPP9, 2:2 and 2:4 DPP9:NLRP1 complexes). One selected 3D class (81,705 particles) was further 3D refined. Subsequently, particles were symmetry expanded (C2) and 3D refined locally followed by per-particle CTFRefine and Bayesian polishing to reach an overall resolution of 3.7 Å for the NLRP1-DPP9 complex. However, the map suffered from anisotropic resolution, and cryoEF^56^ analysis estimated that data collected at a tilt angle of 37° degrees was ideal to fill the orientation gaps in Fourier space.

The overall processing scheme for the 37° tilt data of the NLRP1-DPP9 complex was derived from the workflow described in Relion 3.0^63^. Template-free auto picking with crYOLO^62^ picked 412,419 particles from then pre-processed micrographs. After multiple rounds of 2D classification, 378,410 visually homogenous particles remained for further processing. A random subset comprising 100,000 of these particles was used for *de novo* initial model construction, which was low-passed filtered and used as the initial reference map for 3D classification. After multiple rounds of 3D classification, a 3D class with 89,601 particles remained. The first 3D refinement using these particles yielded an 8.9 Å resolution structure, and the Fourier shell correlation (FSC) curve showed strong fluctuations indicating imprecisions in CTF estimation. To rectify this, we utilized CTF refinement and Bayesian polishing implemented in Relion 3.1, including higher order aberrations correction, anisotropic magnification correction, and per-particle defocus estimation^64^. Iterative rounds of 3D refinement followed by CTF refinement and Bayesian polishing gradually improved the resolution and converged at a resolution plateau of 3.8 Å. Refined particle sets from 0° and 37° stage tilts were then merged and a final 3D refinement was performed. The merged data was reconstructed to an overall resolution of 3.6 Å with much improved FIIND density. Gold-standard FSC between half maps are shown (Extended Data Fig. 1).

A similar data processing scheme was applied to 0° and 37° stage-tilted data of the NLRP1-DPP9-VbP complex. Briefly, crYOLO picked particles were 2D and 3D classified leading to final sets composed of 118,113 and 124,781 particles at 0° and 37° stage-tilts, respectively. 3D refinement followed by one iteration of CTF refinement and Bayesian polishing led to a 3.1 Å resolution map for the non-tilt data. 37° stage-tilted particles initially resolved to 6.1 Å resolution owing to inaccuracies in CTF estimation. To overcome this issue, particles were further refined; iterative 3D refinement followed by CTF refinement (higher order aberrations correction, anisotropic magnification corrections, and per-particle defocus estimation) gradually improved the resolution. Finally, tilted particles were subjected to Bayesian polishing followed by a final round of 3D refinement, which yielded a structure at 3.3 Å resolution. These two refined and polished particle sets (0° and 37° data) were then joined and refined, yielding a final map at 2.9 Å resolution calculated with the gold-standard FSC between half maps (Extended Data Fig. 4).

### Atomic model building

The cryo-EM maps were first fit with the crystal structure of DPP9 dimer (PDB ID: 6EOQ)^48^. A homology model of hNLRP1-FIIND was generated with Schrodinger Prime^65^ using the crystal structure of the rat NLRP1-FIIND (Jijie Chai, personal communications) as a template. Manual adjustment and *de novo* building of missing segments, rigid-body fitting, flexible fitting, and segment-based real-space refinement were performed in distinct parts of the initial model to fit in the density in Coot^66^, with help of UCSF-Chimera^67^ and real-space refinement in Phenix^68^. A few unstructured regions, including parts of the UPA subdomain, were omitted owing to poor density. The full model represents residues Asp18-Met1356 of DPP9, and ZU5 residues Phe1079-Phe1212, UPA^A^ residues Ser1213-Val1350, and UPA^B^ residues Ser1213-Met1356 of NLRP1.

For the VbP-bound structure, the full model represents residues Asp18-Met1356 of DPP9, and ZU5^A^ residues Phe1079-Phe1212, UPA^A^ residues Ser1213-Val1350, and UPA^B^ residues Arg1227-Lys1351 of NLRP1. We modelled covalently linked DPP9 S730-VbP in the cryo-EM map density based on the DPP8-VbP complex structure (PDB ID: 6HP8)^69^ and independently validated fitting with the GemSpot pipeline^70^.

For both structures, interaction analysis was conducted visually and using PISA^71^. Structure representations were generated in ChimeraX^72^, Pymol, and ResMap^73^. Ligand interaction analysis was conducted with Maestro^74^. Pymol and ChimeraX session files are available on our Open Science Framework repository (https://osf.io/x7dv8/). Schematics were created with BioRender.

### CRISPR/Cas9 gene editing

5 × 10^5^ *DPP9* KO HEK293T cells stably expressing Cas9^43^ were seeded in 6-well tissue culture dishes in 2 mL of media per well. The next day cells were transfected according to manufacturer’s instructions (FuGENE HD, Promega) with a mix of four single guide RNA (sgRNA) plasmids targeting DPP8. After 48 h of transfection, cells were transferred to a 10 cm tissue culture dish and selected with puromycin (2 μg/mL) until untransfected control cells were dead. Single cell clones were isolated by serial dilution and confirmed by genomic sequencing (Extended Data Fig. 7b).

### Immunoblotting

Samples were run on either NuPAGE™ 4 to 12%, Bis-Tris 1.0 mm, Mini Protein Gel (Invitrogen) for 30 min at 175 V or NuPAGE™ 4 to 12%, Bis-Tris, 1.0 mm, Midi Protein Gel (Invitrogen) for 45-60 min at 175 V. Gels were transferred to nitrocellulose with the Trans-Blot Turbo Transfer System (BIO-RAD). Membranes were blocked with Intercept™ (TBS) Blocking Buffer (LI-COR) for 30 min at ambient temperature, prior to incubating with primary antibody overnight at 4 °C. Blots were washed 3 times with TBST buffer prior to incubating with secondary antibody for 60 min at ambient temperature. Blots were washed 3 times, rinsed with water and imaged via Odyssey CLx (LI-COR). Primary antibodies used in this study include: DPP9 rabbit polyclonal Ab (Abcam, Ab42080), FLAG® M2 mouse monoclonal Ab (Sigma, F3165), GAPDH rabbit monoclonal Ab (Cell Signaling Technology, 14C10), NLRP1 mouse monoclonal Ab (R&D Systems, MAB6788), CASP1 rabbit polyclonal Ab (Cell Signaling Technology, 2225), GSDMDC1 rabbit polyclonal Ab (Novus, NBP2-33422), V5 rabbit polyclonal Ab (Abcam, Ab9116). Secondary antibodies used in this study include: IRDye 680 RD Streptavidin (LI-COR, 926-68079), IRDye 800CW anti-rabbit (LI-COR, 925-32211), IRDye 800CW anti-mouse (LI-COR, 925-32210), IRDye 680CW anti-rabbit (LI-COR, 925-68073), IRDye 680CW anti-mouse (LI-COR, 925-68072).

### Immunoprecipitation assays

HEK293T cells were seeded at 5 × 10^5^ cells/well in 6-well tissue culture dishes. The following day the cells were transfected with plasmids encoding the indicated FLAG-tagged protein (2 μg) with FuGENE HD according to manufacturer’s instructions (Promega). After 48 h cells were harvested and washed 3 times with PBS. Pellets were lysed in Tris-Buffered Saline (TBS) with 0.5% NP-40 using pulse sonication and centrifuged at 20,000 x *g* for 10 min at 4 °C. Protein concentration of the soluble proteome was determined using the DC Protein Assay kit (Bio-Rad) and adjusted to 2 mg/mL. Lysates were treated with DMSO or VbP (10 µM) for 1 h. Lysates were incubated with 20 µL of anti-FLAG-M2 agarose resin (Sigma) overnight at 4 °C. After washing 3 × 500 µL with cold PBS in microcentrifuge spin columns (Pierce), bound proteins were eluted by incubating resin with 40 µL of PBS with 150 ng/μL 3x-FLAG peptide for 1 h at 4 °C. An equal volume of 2x sample loading was added to the eluate and boiled. Immunoblots were developed with the Odyssey CLx imaging system (LI-COR).

For DPP9 mutants, *DPP8/9* double knockout (DKO) HEK293T cells were seeded at 1 × 10^6^ cells/well in 6-well tissue culture dishes. The following day cells were transfected with plasmids encoding for FLAG-tagged NLRP1 (2 μg) or the indicated DPP9 construct (2 µg) with FuGENE HD according to manufacturer’s instructions (Promega). Cells were harvested, lysed, and normalized as per above. Lysates were mixed in a one to one ratio of DPP9-NLRP1 prior to treating with DMSO or VbP (10 µM) for 1 h. Assay was completed as above.

### DPP9 activity assay

A solution of substrate (1 mM Gly-Pro-AMC) was prepared in DMSO. 19 µL of PBS, recombinant DPP9 (1 nM), or the indicated cell lysate was added to a 384-well, black, clear-bottom plate (Corning) followed by 1 µL of substrate to initiate the reaction. Substrate cleavage was measured as increasing fluorescence signal recorded at ambient temperature every minute at 380-nm excitation and 460-nm emission wavelengths over a 30 min period. Activity was calculated by linear regression (Prism 7). For NLRP1 peptide assays, peptide or DMSO was added to the indicated final concentration and incubated for 30 min on ice prior to addition of GP-AMC substrate.

### LDH cytotoxicity assay

HEK293T cells stably expressing CASP1 and GSDMD were seeded at 1 × 10^5^ cells/well in 12-well tissue culture dishes. The following day the cells were transfected with plasmids encoding for ASC (0.01 µg), the indicated NLRP1 construct (0.02 μg) and RFP (to 1 µg) with FuGENE HD according to manufacturer’s instructions (Promega). At 24 h cells were treated with DMSO or VbP (10 µM). 24 h later supernatants were analysed for LDH activity using the Pierce LDH Cytotoxicity Assay Kit (ThermoFisher) and lysate protein content was evaluated by immunoblotting. For NLRP1-CT experiments LDH was measured and lysates were collected 24 h post transfection.

### ASC puncta formation assay

HEK293T cells were seeded at 2.5 × 10^5^ cells/mL in 12-well tissue culture dishes. The following day the cells were transfected with a master mix (50 µL per well) of plasmids encoding for GFP-ASC (0.033 µg), the indicated NLRP1 construct (0.066 μg), and RFP (to 2 µg) with FuGENE HD according to manufacturer’s instructions (Promega). After 24 h cells were treated with DMSO or VbP (10 µM) as indicated for 6 h. Nuclei were stained by treating cells with Hoechst dye (0.1 µL of 10 mg/ml solution). Live cell imaging was performed on a Zeiss Axio Observer.Z1 inverted wide-field microscope using 40×/0.95NA air objective. For each well, 15 positions were imaged on DAPI, red, and green fluorescence channels at a single time point from a given experiment. Data were exported as raw .czi files and analyzed using custom macro written in ImageJ/FIJI. Total cell area was estimated from RFP-positive signal, and the number of GFP-ASC specks was quantified using the “Analyze particles” function following threshold adjustment in the GFP positive images.

### Chemical enrichment of protease substrates (CHOPS)

*DPP8/9* DKO HEK293T cells were seeded at 1 × 10^6^ cells/well in 6-well tissue culture dishes. The following day cells were transfected with plasmids encoding for FLAG-tagged NLRP1 (2 μg) with FuGENE HD according to manufacturer’s instructions (Promega). After 48 h, cells were harvested, washed 3x with PBS, lysed in PBS with pulse sonication, and centrifuged at 20,000 x *g* for 10 min at 4 °C. Protein concentration of the soluble proteome was determined using the DC Protein Assay kit (Bio-Rad) and adjusted to 1.5 mg/mL.

250 µL of lysate was incubated with either PBS or recombinant DPP9 (rDPP9, ∼3 µg). The activity of rDPP9 was confirmed by GP-AMC assay prior to its addition. The mixtures were allowed to incubate for 16 h at 37 °C, after which samples were boiled for 10 min to deactivate rDPP9. 2PCA-biotin probe was added to lysates (10 mM final concentration) and incubated with shaking at 37 °C for an additional 16 h. Excess probe was removed by buffer exchange into fresh PBS with Amicon Ultra-0.5 mL Centrifugal Filters, 10k MW cutoff (3 × 500 µL). SDS was added to 1% and samples were boiled for 10 min, prior to diluting further with 5 mL of PBS. 100 µL of high capacity neutravidin agarose resin (Pierce) was added, before rotating tubes end-over-end for 1 h at room temperature. Samples were centrifuged at 1,000 x *g* for 2 min, supernatant was removed and beads were washed with 10 mL of PBS. This was step was repeated for a total of 3 times. Bound proteins were eluted by boiling the resulting neutravidin resin in 100 µL of 2x loading dye and evaluated by immunoblot.

### Data availability

The cryo-EM maps have been deposited in the Electron Microscopy Data Bank under the accession numbers EMD-22074 (NLRP1-DPP9) and EMD-22075 (NLRP1-DPP9-VbP). The atomic coordinates have been deposited in the Protein Data Bank under the accession numbers 6×6A (NLRP1-DPP9) and 6×6C (NLRP1-DPP9-VbP). Raw cryo-EM data will be deposited into EMPIAR. Extended protein purification protocols will be made available on Protocols.io (https://www.protocols.io/groups/hao-wu-lab). Pymol session files and full-sized figures will be made available on OSF (https://osf.io/x7dv8/). All constructs will be made available on Addgene. All other data can be obtained from the corresponding authors upon reasonable request.

## Acknowledgements

We thank Dr. Tian-min Fu of the Wu lab for initiating an apo-NLRP1 project and all members of the Wu and Bachovchin labs for helpful discussions. We thank Dr. Jijie Chai for sharing their crystal structure of the rat NLRP1 FIIND prior to submission, Dr. Min Su for his advice on running goCTF and CTF estimation for tilt datasets, Dr. Maria Ericsson at the HMS EM facility for advice and training, and Drs. Richard Walsh, Sarah Sterling, and Shaun Rawson at the Harvard Cryo-EM Center for Structural Biology for cryo-EM training and productive consultation. We also thank Drs. Georgios Skiniotis and Mike Robertson for sharing their GemSpot script and for troubleshooting advice. We thank Dr. Thomas Edwards and the National Cancer Institute’s National Cryo-EM Facility at the Frederick National Laboratory for Cancer Research under contract HSSN261200800001E. We also thank Dr. Janette Myers and the Pacific Northwest Center for Cryo-EM at Oregon Health & Science University for screening and preliminary dataset collection, under the NIH grant U24GM129547 and accessed through EMSL (grid.436923.9), a DOE Office of Science User Facility sponsored by the Office of Biological and Environmental Research. The pcDNA3.1 LIC 6A and pcDNA 3.1 LIC 6D plasmids were gifts from Scott Gradia (Addgene plasmids #30124 and #30127). Our research used software and computing support at SBGrid and the Extreme Science and Engineering Discovery Environment (XSEDE) Bridges at the Pittsburgh Supercomputing Center through allocation TG-MCB190086. This work was supported by National Institutes of Health (DP1 HD087988 and R01 AI124491 to H.W., T32 GM007739-Andersen and F30 CA243444 to A.R.G., and the Memorial Sloan Kettering Cancer Center Core Grant P30 CA008748 and R01 AI137168 to D.A.B.).

## Author contributions

L.R.H., H.S. and H.W. conceived the study. L.R.H. designed constructs with input from H.S. L.R.H., H.S., P.F., and K.D. carried out preliminary expression and purification studies. L.R.H. purified the complexes. H.S. and L.R.H. made cryo-EM grids for data collection. L.R.H. screened cryo-EM grids and collected cryo-EM data. H.S. and L.R.H. analysed cryo-EM data. H.S. performed model building and refinement. L.R.H. and H.S. designed mutants for in vitro and cell-based assays. A.R.G. performed all cell-based assays, peptide mass spectrometry, and CHOPS analysis under D.A.B.’s supervision. J.M. and J.P. performed crosslinking mass spectrometry and data analysis under S.P.G.’s supervision. A.R.G., L.R.H., H.S., H.W., and D.A.B. designed the experiments. L.R.H, H.S. and H.W. wrote the manuscript with input from other authors.

---

**Extended Data Fig. 1.**
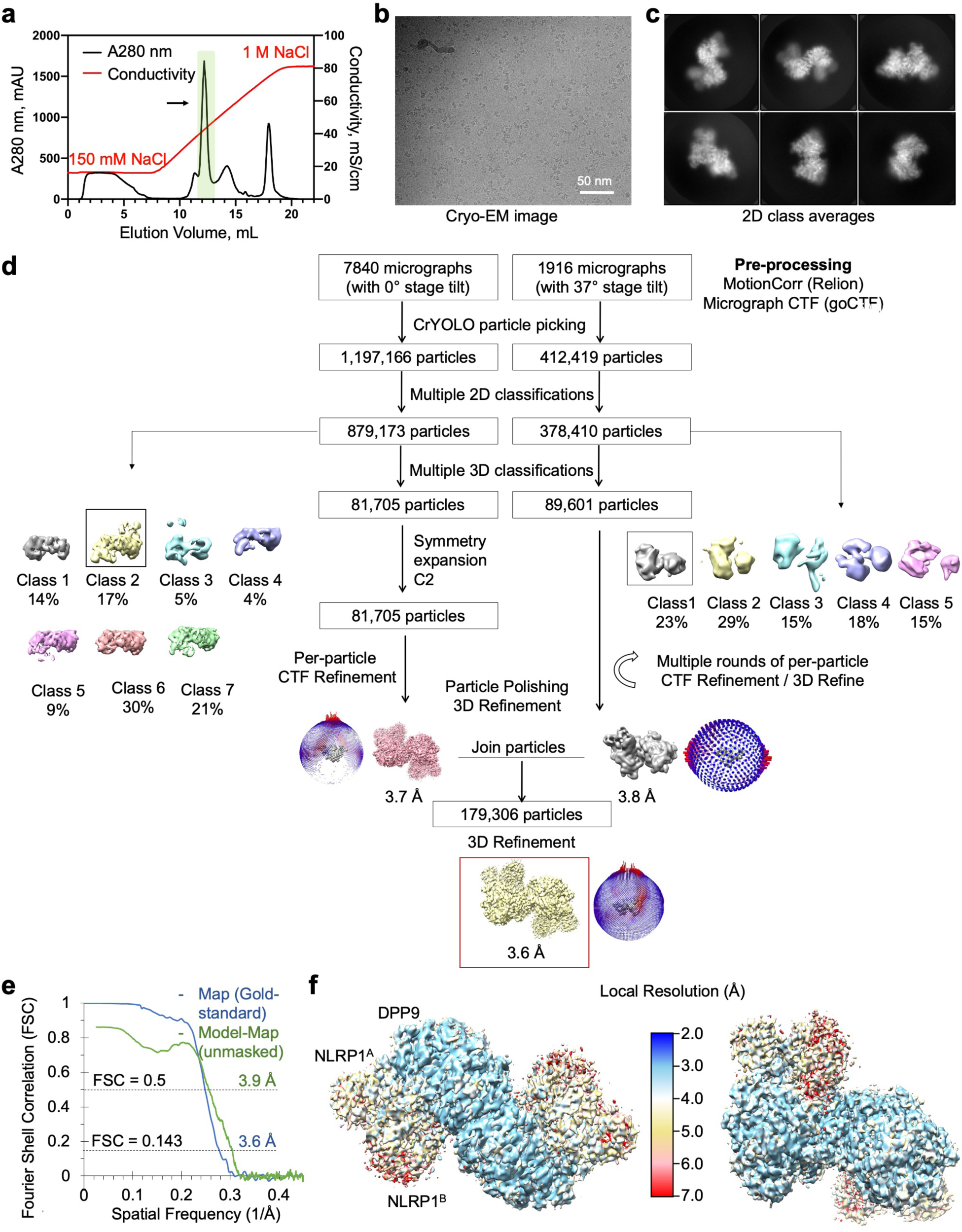
Structure determination of the NLRP1-DPP9 complex. **a**, Purification of the NLRP1-DPP9 complex by ion exchange chromatography. The ternary complex peak is shaded in green and labelled with an arrow. **b**, A representative cryo-EM micrograph. **c**, Representative 2D class averages. **d**, Workflow for the NLRP1-DPP9 complex structure determination. **e**, Map-map and map-model FSC curves. **f**, Local resolution distribution of the final map calculated with ResMap^73^.

**Extended Data Fig. 2.**
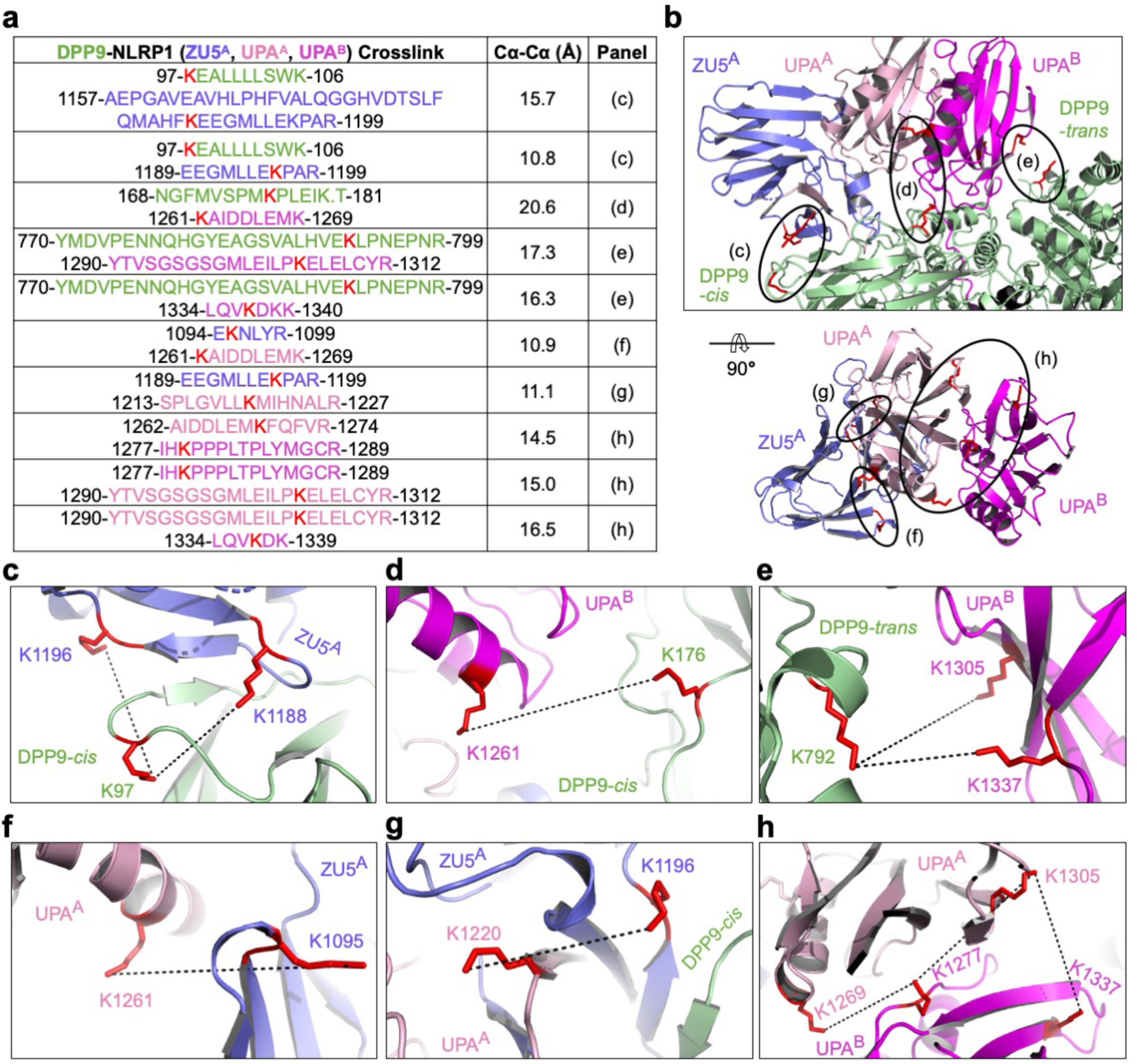
Crosslinking mass spectrometry analysis of the NLRP1-DPP9 complex. **a**, Summary of BS3 crosslinking between DPP9 and NLRP1. High confidence crosslinked peptides are displayed, with residue ranges labelled and colours coded by domains. Crosslinked lysine pairs are indicated in red. Cα-Cα distances between these lysine residues interpreted by the final NLRP1-DPP9 model are shown, along with the figure panel names for their detailed depictions. **b**, Overview of BS3-mediated crosslinks. **c-h**, Zoom-ins highlighting crosslinked lysine pairs (red) interpreted by the final NLRP1-DPP9 model.

**Extended Data Fig. 3.**
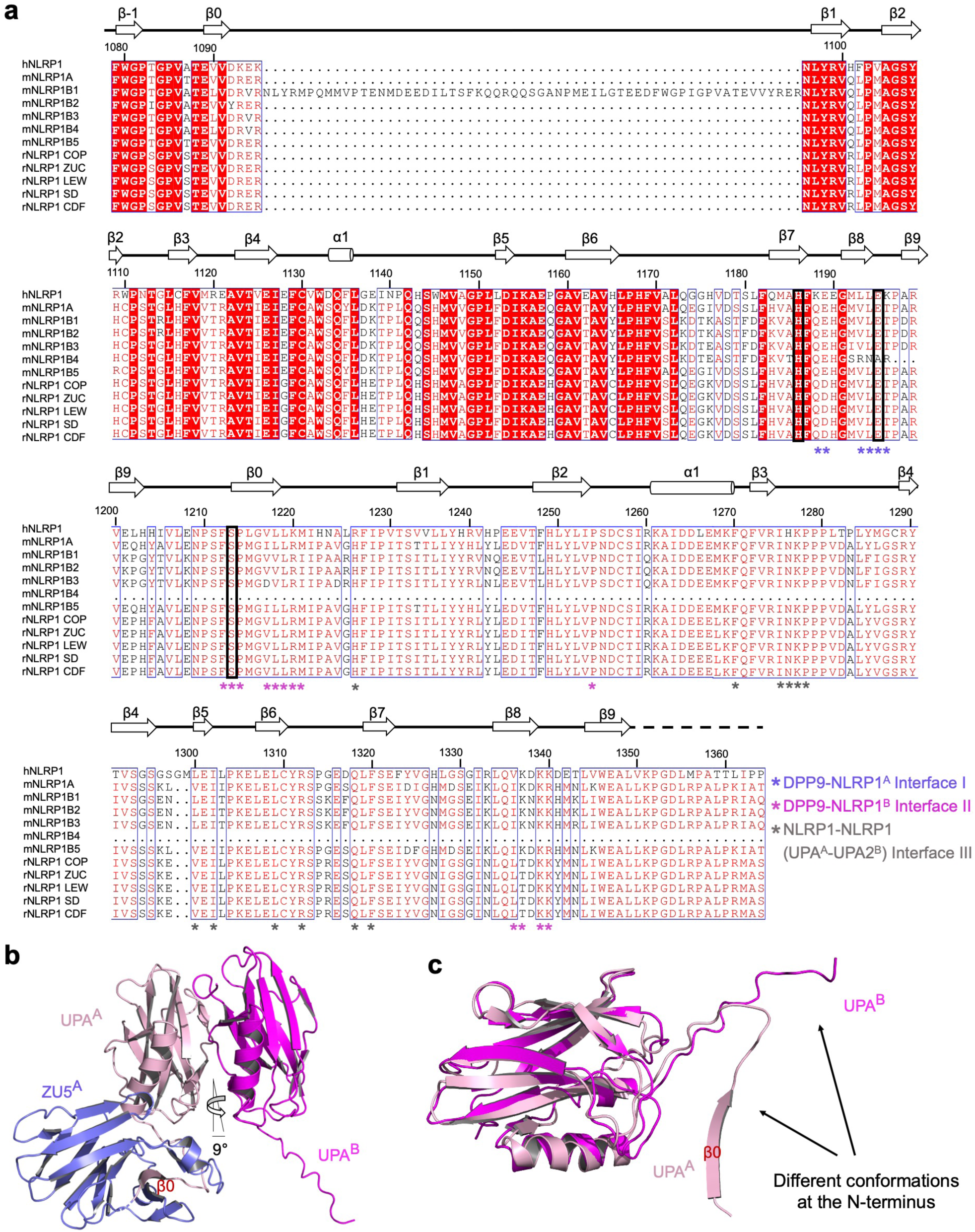
Sequence and structural analysis of FIIND. **a**, ClustalW multiple sequence alignment between human NLRP1 (hNLRP1), mouse NLRP1 (mNLRP1, different isoforms), and rat NLRP1 (rNLRP1, different isoforms). COP: Copenhagen; ZUC: Zucker; LEW: Lewis; SD: Sprague Dawley; and CDF: Fischer. Secondary structures and residue numbers are denoted based on the human FIIND^A^ structure in the NLRP1-DPP9 ternary complex. Interfacial residues in the NLRP1-DPP9 complex are annotated with asterisks, and residues in the catalytic triad (H1186, E1195, S1213) are boxed in black. **b**, The ZU5^A^-UPA^A^-UPA^B^ module that binds DPP9. UPA^A^ and UPA^B^ interact with each other in a front-to-back manner with only a 9° rotation between them. **c**, Altered conformation of the UPA^B^ N-terminus that binds in the DPP9 active site tunnel in comparison to UPA^A^ in a complete FIIND^A^.

**Extended Data Fig. 4.**
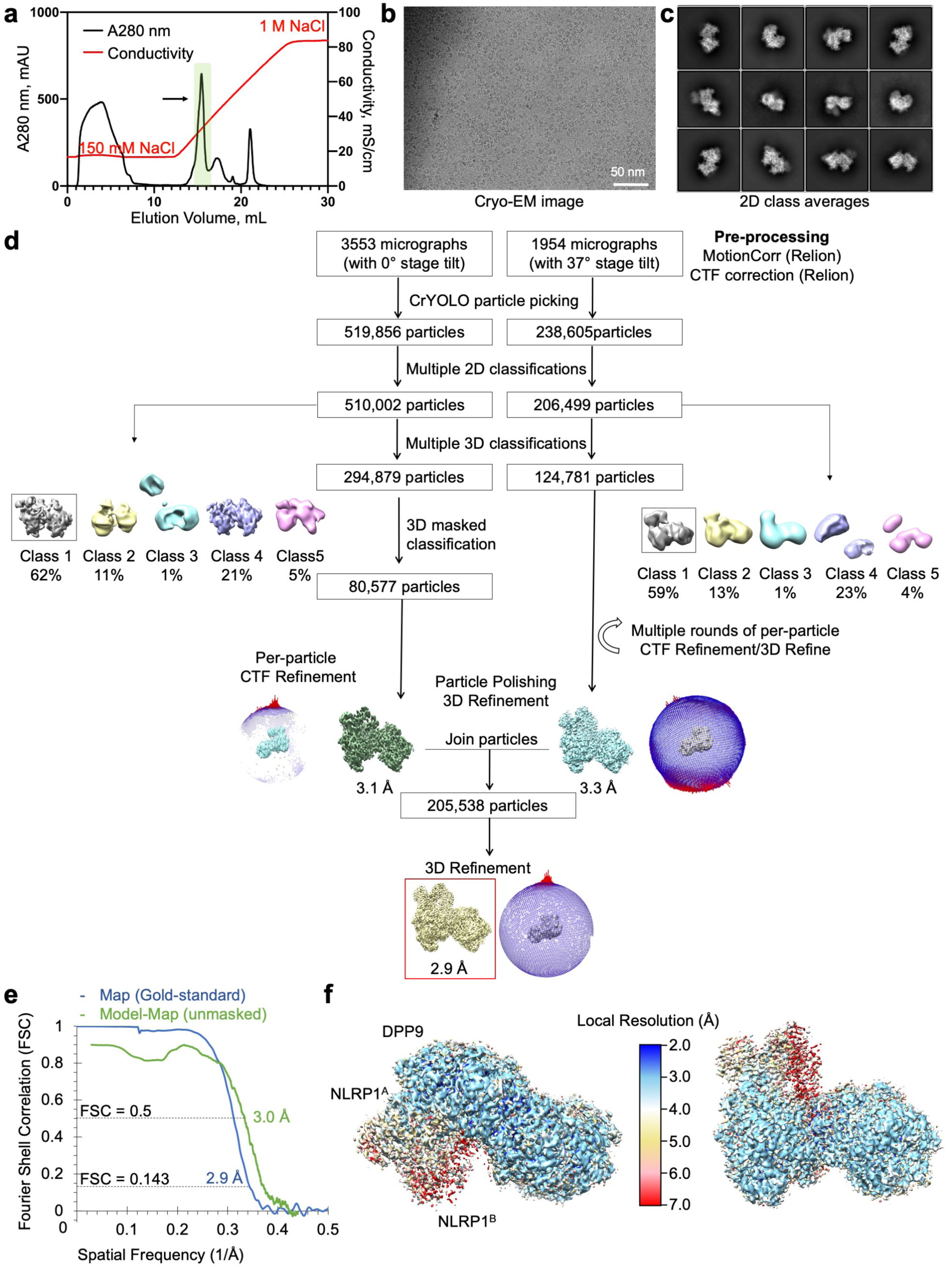
Structure determination of the NLRP1-DPP9 complex with VbP. **a**, Purification of the NLRP1-DPP9 complex in the presence of VbP by ion exchange chromatography. The ternary complex peak is shaded in green and labelled with an arrow. **b**, A representative cryo-EM micrograph. **c**, Representative 2D class averages. **d**, Workflow for the NLRP1-DPP9-VbP complex structure determination. **e**, Map-map and map-model FSC curves. **f**, Local resolution distribution of the final map calculated with ResMap^73^.

**Extended Data Fig. 5.**
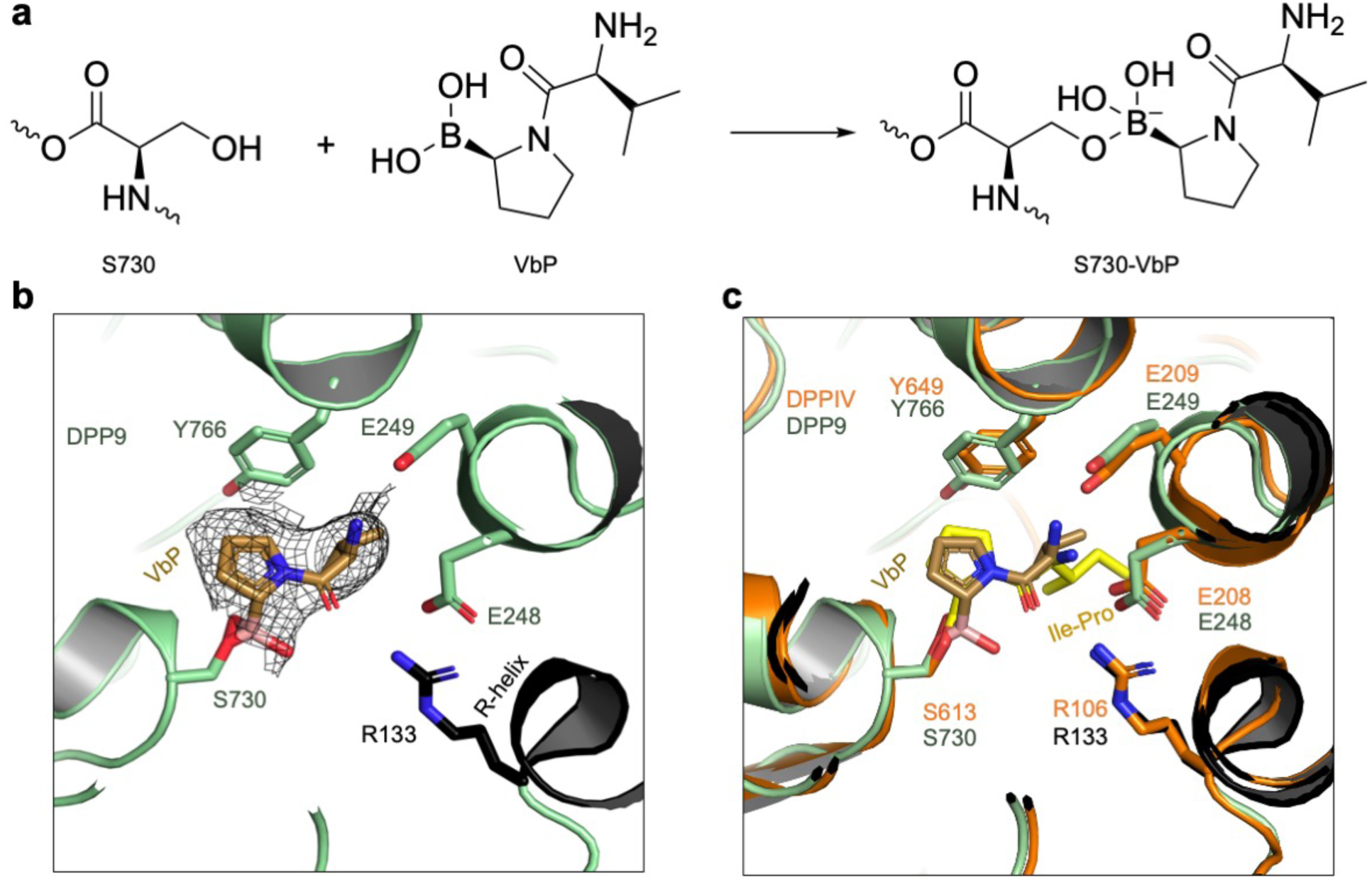
VbP interactions in the DPP9 active site and comparison to a DPP substrate. **a**, Schematic of covalent linkage between DPP9’s S730 and VbP. **b**, Fit of VbP into the cryo-EM density. VbP is shown in stick with carbon atoms in light brown. The charged amino group of VbP interacts with the DPP9 EE loop which also coordinates a substrate N-terminus, and the carbonyl oxygen of VbP interacts with R133 of the R-helix. The covalent linkage of VbP with S730, the catalytic serine, is displayed. **c**, Structural alignment of the VbP-bound DPP9 model (green) and the crystal structure of bacterial DPP4 bound to the substrate Ile-Pro (PDB ID: 5YP3, orange)^49^. VbP assumes a pose remarkably like a model substrate.

**Extended Data Fig. 6.**
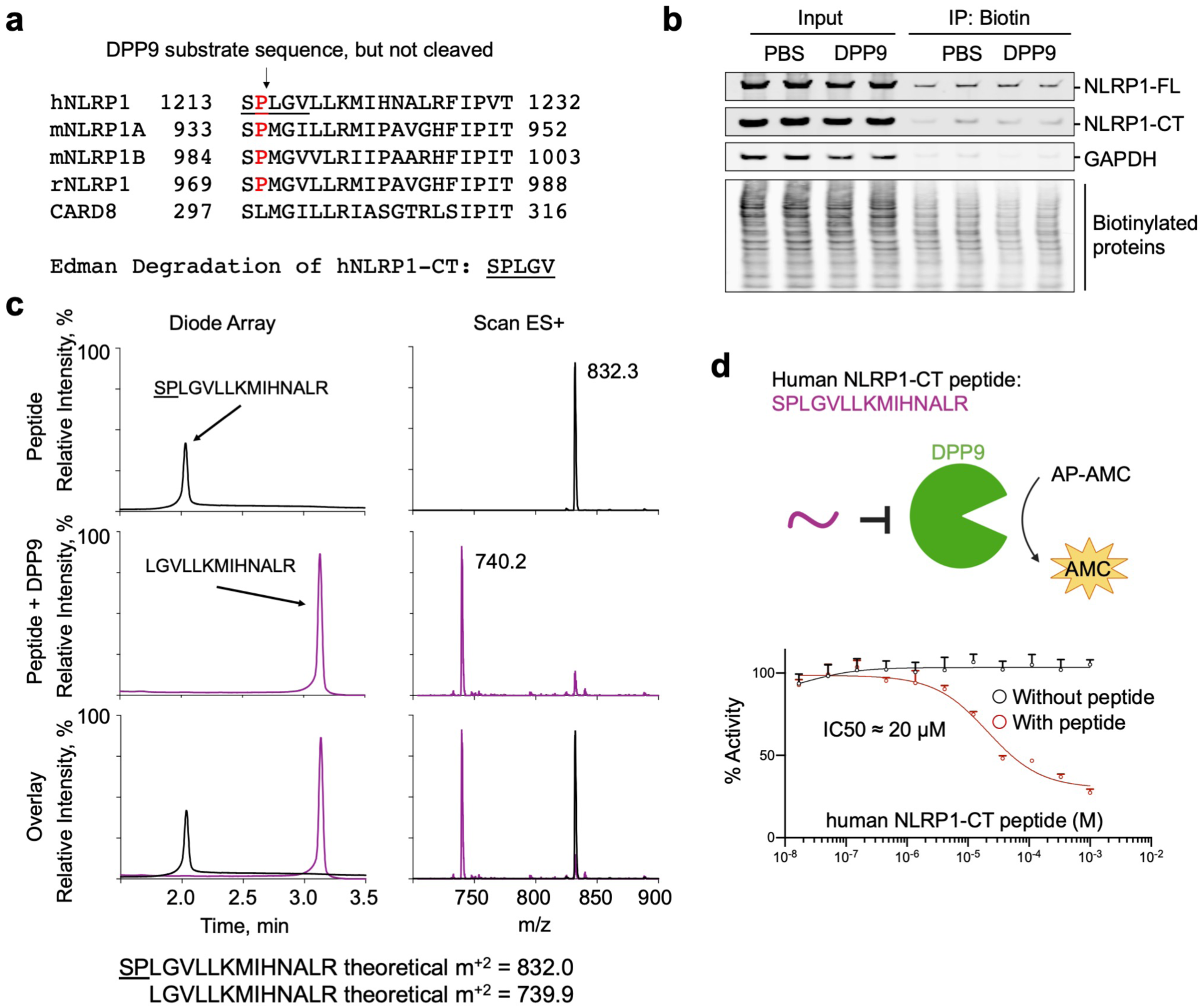
Lack of cleavage of intact NLRP1-CT but the cleavage of its isolated N-terminal peptide by DPP9. **a**, N-terminal sequencing of the purified NLRP1-DPP9 complex showing that the NLRP1-CT is not cleaved by the co-expressed DPP9. **b**, CHOPS assay showing that DPP9 does not cleave NLRP1-CT. Briefly, WT NLRP1 expressed in *DPP8/9* DKO HEK293T cells was incubated with PBS or recombinant DPP9 prior to labelling with the 2PCA biotin probe to capture biotinylated proteins. The inputs and the eluents were analysed by immunoblots using anti-NLRP1-CT (NLRP1-FL and NLRP1-CT), anti-GAPDH, and anti-Biotin antibodies. **c**, Evidence of cleavage of the isolated 15 residue N-terminal peptide in NLRP1-CT by recombinant DPP9 from mass spectrometry analysis. **d**, Inhibition of DPP9 catalytic activity against AP-AMC by the isolated NLRP1-CT peptide. All data are representative of > 2 independent experiments.

**Extended Data Fig. 7.**
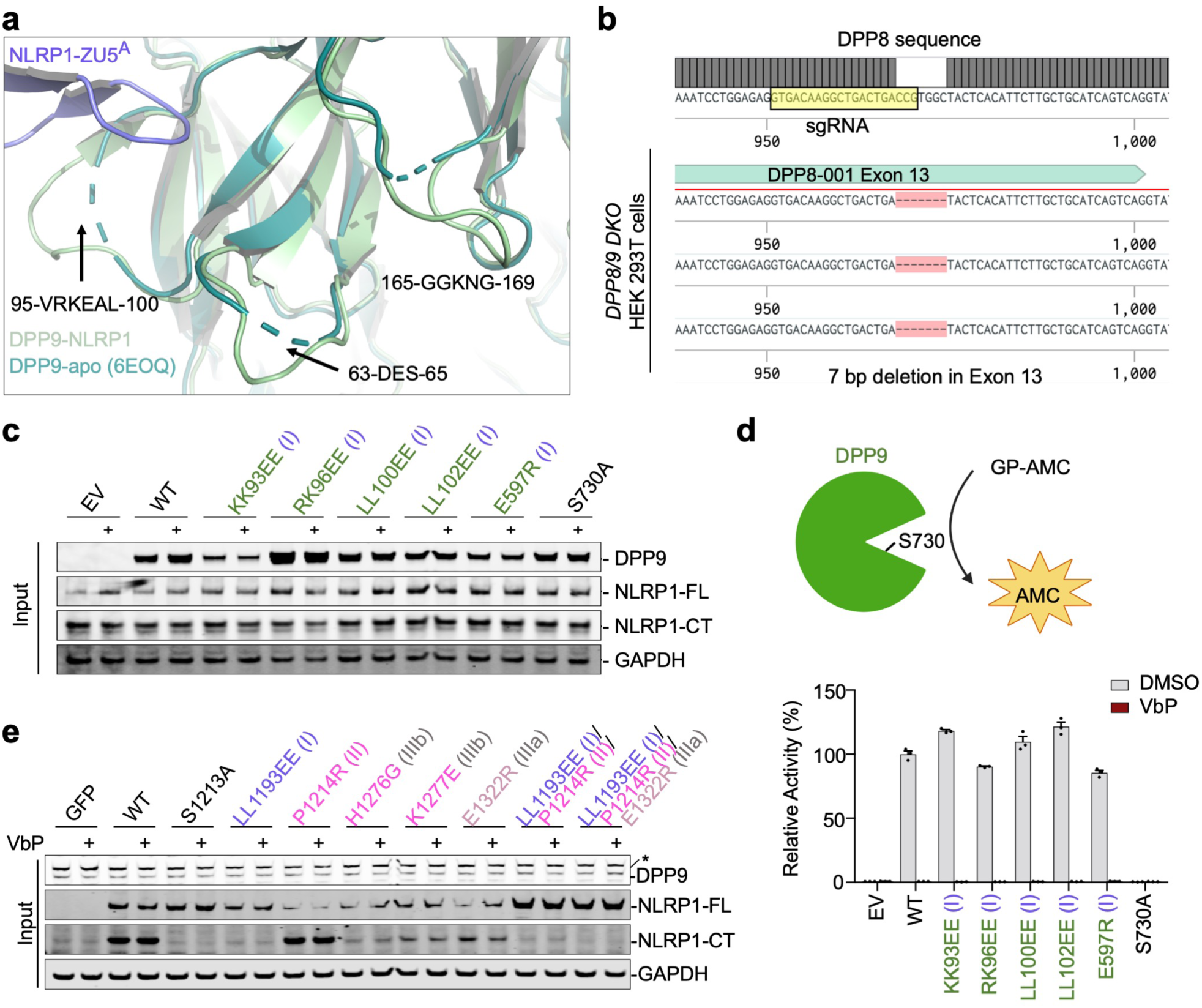
Mutational analysis of the interactions in the NLRP1-DPP9 ternary complex. **a**, Disorder to order transition of several DPP9 surface loops from the isolated DPP9 crystal structure (PDB ID: 6EOQ)^48^ to the NLRP1-bound DPP9 cryo-EM structure. **b**, Genomic confirmation of DPP8 KO in *DPP8/9* DKO HEK293T cells. Single guide RNA (sgRNA) sequence is highlighted. **c**, Immunoblots of the input lysates for the FLAG co-immunoprecipitation with WT or mutant DPP9 and WT NLRP1-FLAG. **d**, Cleavage rate of a model DPP9 substrate, GP-AMC, by WT DPP9 and its structure-guided mutants. Only the catalytically dead mutant S730A disrupts catalytic activity and sensitivity to the DPP9 inhibitor, VbP. **e**, Immunoblots of the input lysates for the FLAG co-immunoprecipitation with WT or mutant NLRP1-FLAG and WT DPP9. All data are representative of > 2 independent experiments.

**Extended Data Fig. 8.**
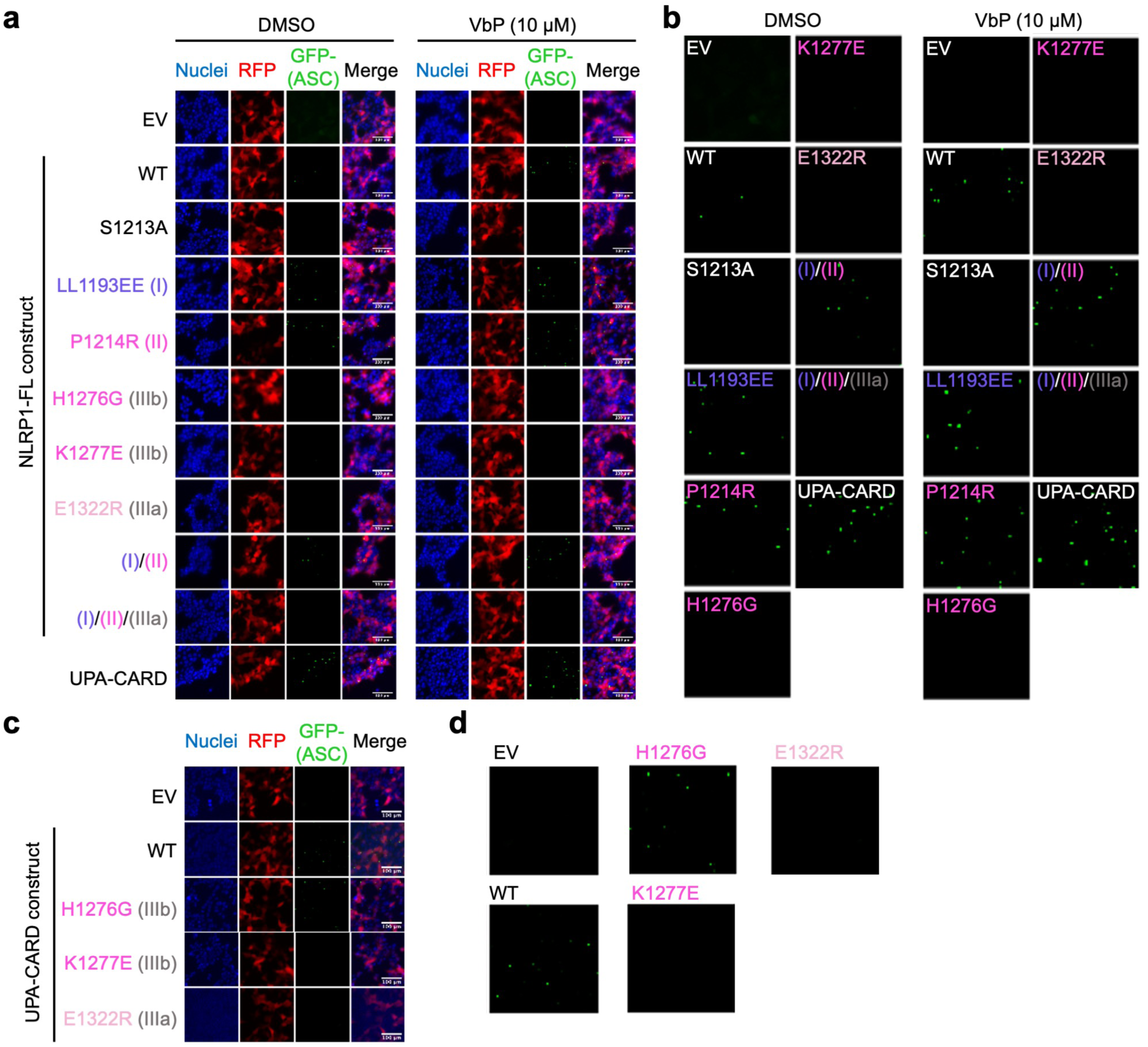
Confocal microscopy of ASC specks. **a**, Co-expression of GFP-ASC and NLRP1-FL (WT and mutants) or UPA-CARD (positive control). Autoactive constructs form GFP-ASC specks in the absence of VbP. Image quantitation is shown in Fig. 3b. **b**, Enlarged GFP-ASC panels from (a). **c**, Co-expression of GFP-ASC and UPA-CARD (WT and interface III mutants). Image quantitation is shown in Fig. 3d. **d**, Enlarged GFP-ASC panels from (c). All data are representative of > 3 independent experiments.

**Extended Data Fig. 9.**
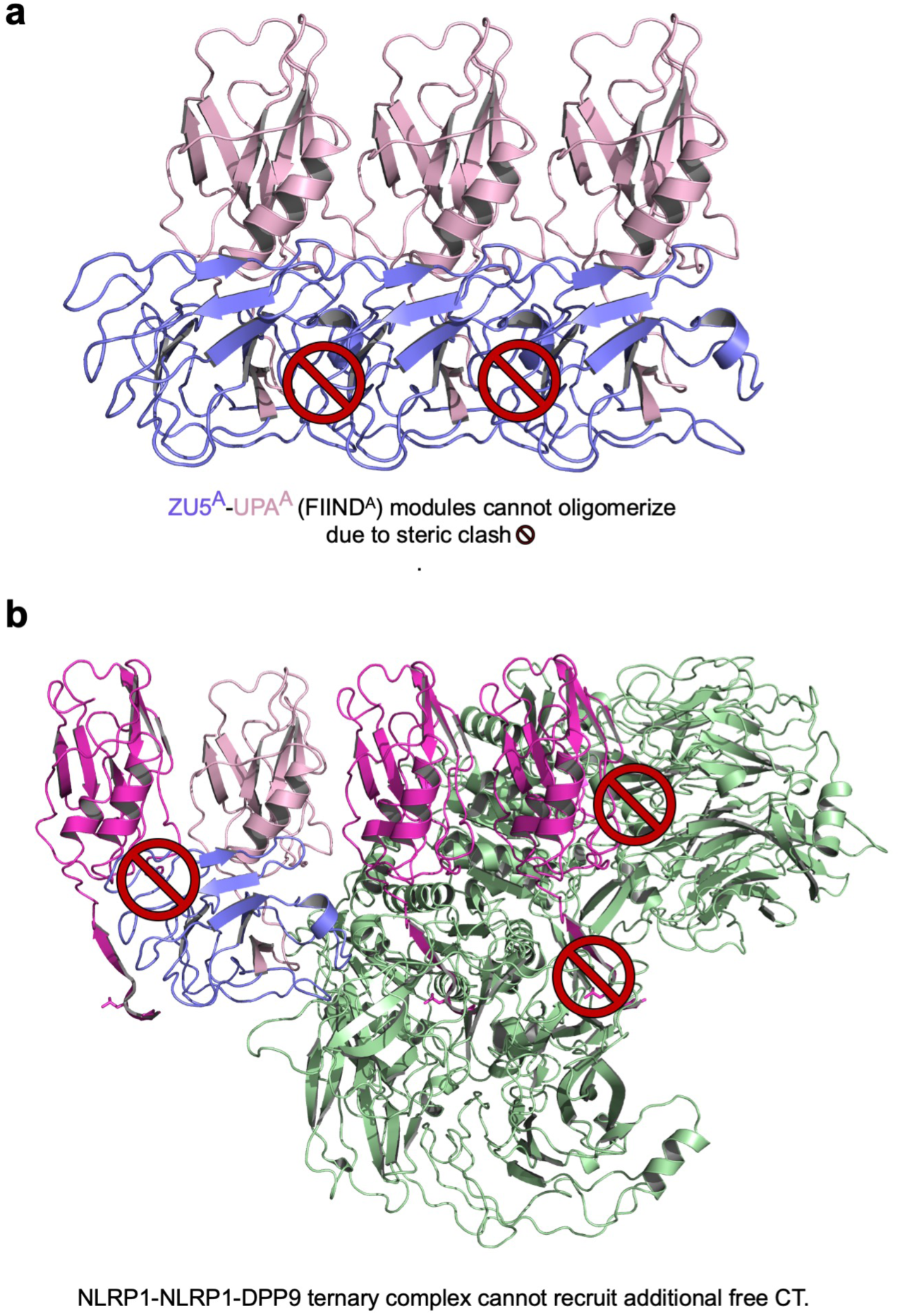
The ZU5 domain and DPP9 sterically hinder UPA polymerization. **a**, Modelling of a FIIND polymer using the observed UPA^A^-UPA^B^ relationship. Adjacent ZU5 molecules would clash, suggesting that UPA polymerization can only occur for free NLRP1-CTs. **b**, Modelled recruitment of free UPA at both UPA^A^ and UPA^B^ in the ternary complex with DPP9. The additional UPA subdomain next to FIND^A^ clashes with the ZU5 subdomain, and the additional UPA next to UPA^B^ clashes with both DPP9 monomers in the complex, suggesting that DPP9 inhibits UPA oligomerization.

**Extended Data Table 1.** Cryo-EM data collection and refinement statistics of NLRP1-DPP9

|  | **NLRP1-DPP9** |  | **NLRP1-DPP9-VbP** |  |
| --- | --- | --- | --- | --- |
| Stage tilt (°) | 0° | 37° | 0° | 37° |
| Movies collected | 7,840 | 1,916 | 3,553 | 1,954 |
| Magnification | 105,000 | 105,000 | 105,000 | 105,000 |
| Voltage (kV) | 300 | 300 | 300 | 300 |
| Electron exposure (e<sup>-</sup>/Å<sup>2</sup>) | 67.54 | 67.6 | 52 | 65 |
| Defocus range (µm) | -1.0 to -2.5 | -1.5 to -3.0 | -0.8 to -2.2 | -1.5 to -3.0 |
| Exposure time (s) | 1.8 | 2.6 | 2.0 | 2.4 |
| Pixel size (Å) | 0.825 | 0.825 | 0.825 | 0.825 |
| Symmetry imposed | C1 | C1 | C1 | C1 |
| Initial particle images (no.) | 758,461 | 398,698 | 1,1609,585 | 239,416 |
|  | **Merged Data** |  | **Merged Data** |  |
| Final particle images (no.) | 179,306 |  | 205,538 |  |
| Map resolution (Å) | 3.6 |  | 2.9 |  |
| FSC threshold | 0.143 |  | 0.143 |  |
| **Refinement** |  |  |  |  |
| Model resolution (Å) | 3.9 |  | 2.9 |  |
| FSC threshold | 0.5 |  | 0.5 |  |
| Map sharpening *B* factor (Å<sup>2</sup>) | -81.4 |  | -49.0 |  |
| **Model composition** |  |  |  |  |
| Non-hydrogen atoms | 20,269 |  | 16,741 |  |
| Protein residues | 2,524 |  | 2,080 |  |
| Ligands |  |  | 2 |  |
| B factors (Å<sup>2</sup>) | 110.1 |  | 82.7 |  |
| **R.m.s. deviations** |  |  |  |  |
| Bond lengths (Å) | 0.017 |  | 0.008 |  |
| Bond angles (°) | 1.633 |  | 1.363 |  |
| **Validation** |  |  |  |  |
| MolProbity score | 1.77 |  | 1.88 |  |
| Clashscore | 5.59 |  | 7.21 |  |
| Poor rotamers (%) | 0.00 |  | 0.00 |  |
| **Ramachandran plot** |  |  |  |  |
| Favoured (%) | 92.6 |  | 93.3 |  |
| Allowed (%) | 7.2 |  | 6.5 |  |
| Disallowed (%) | 0.2 |  | 0.2 |  |

